# AKAP-CaN-NFATc3 signaling drives persistent inhibitory long-term potentiation

**DOI:** 10.64898/2026.11.10.765705

**Authors:** Sarah E. Wolfe, Katlin H. Zent, Joshua D. Garcia, Sara E. Gookin, Mark L. Dell’Acqua, Katharine R. Smith

## Abstract

GABAergic inhibitory synaptic plasticity has critical roles in neuronal function, synaptic plasticity, and cognition. Activity-dependent alterations in gene expression are essential for generating long-term changes in inhibitory postsynaptic strength, yet the signaling mechanisms that drive inhibitory postsynaptic plasticity are not well understood. During an important form of inhibitory long-term potentiation (iLTP), NMDA receptor (NMDAR) activity promotes *de novo* synthesis of GABA_A_ receptors (GABA_A_Rs) and the inhibitory scaffold, gephyrin, leading to persistent strengthening of inhibition. However, it remains unclear what activity-dependent signaling mechanisms control these changes in gene expression to maintain iLTP. In this study, we identify that activation of the transcription factor, nuclear-factor of activated T-cells (NFAT), and anchoring of the phosphatase, calcineurin (CaN), to A-kinase anchoring protein (AKAP79/150) are crucial in regulating long-term structural and functional GABAergic postsynaptic plasticity. Using short hairpin RNA (shRNA) to knock-down hippocampal NFAT isoform, NFATc3, we demonstrate its requirement for the synthesis of new inhibitory synaptic proteins, inhibitory synapse growth, and strengthening of synaptic currents during persistent iLTP. Additionally, using a knock-in mouse line containing a mutation in the CaN binding region of AKAP, we find that AKAP-CaN signaling lies upstream of NFATc3 activation and is required for increased inhibitory synapse clustering and synapse strengthening during iLTP via regulation of miRNA transcriptional activity and protein synthesis. Thus, we have identified an excitation-transcription coupling pathway that rapidly controls the synthesis of new inhibitory synaptic proteins and maintains persistent inhibitory synaptic plasticity, an essential process for memory formation and cognition in the brain.

## INTRODUCTION

Long-term synaptic plasticity is the cellular underpinning of learning and memory, requiring synthesis of new proteins to induce sustained changes in excitation and inhibition (1–4). *De novo* protein synthesis during plasticity stabilizes and strengthens pre- and postsynaptic structures leading to sustained modulation of synaptic strength (5–9). Alterations in protein expression during synaptic plasticity require modifications in transcription in response to neuronal excitation, known as excitation-transcription (E-T) coupling (10). E-T coupling mechanisms have been well characterized in the context of high-activity stimuli inducing strong calcium (Ca^2+^) influx, leading to processes such as excitatory long-term potentiation (eLTP) (10), yet, the E-T coupling mechanisms that underly long-term changes in GABAergic inhibitory synaptic function are poorly understood.

Inhibitory synaptic plasticity is critical for modulating excitability, plasticity, and circuit function (11). An important form of inhibitory postsynaptic plasticity, known as inhibitory long-term potentiation (iLTP), is initiated by moderate NMDA receptor (NMDAR) activation leading to the strengthening of GABAergic inhibitory synapses (12–16). Within 20 minutes following iLTP induction, trafficking of pre-existing GABA_A_ receptors (GABA_A_Rs) and the inhibitory synaptic scaffold, gephyrin, to inhibitory synapses increases GABAergic synaptic activity (7, 14–16). Over longer timescales (45-90 minutes following NMDAR activation), iLTP requires *de novo* synthesis of GABA_A_R subunits and gephyrin to sustain increased inhibitory synaptic strength (6, 7). We previously identified two short non-coding microRNAs (miRNAs), miR376c and miR153, that target and repress the translation of the mRNAs encoding synaptic GABA_A_R subunits (GABA_A_R- α1 and GABA_A_R-γ2) and gephyrin, respectively. Upon iLTP induction, transcription of these miRNAs decreases, which consequently permits the translation of GABA_A_R subunits and gephyrin mRNAs, leading to the sustained strengthening of GABAergic synapses (6, 7).

Additionally, we identified active binding sites for the transcription factor, nuclear-factor of activated T-cells (NFATc3) within upstream regulatory regions of miR376c and miR153 genes, and *in vitro* assays suggest that NFATc3 promotes transcriptional repression of these miRNAs in response to iLTP stimulation (6, 7). However, whether NFATc3 is required for iLTP and what upstream signaling mechanisms direct its actions remains untested.

In response to high-activity levels, cytosolic NFATc3 rapidly translocates into the nucleus following dephosphorylation by calcineurin (CaN), via activation of L-type calcium channels (LTCCs) (17–19). We previously observed *de novo* protein synthesis during iLTP requires the activation of CaN, as well as LTCC-dependent Ca^2+^ influx (6, 7), suggesting that a similar CaN-NFAT-dependent signaling mechanism could regulate transcription during iLTP. Furthermore, NFAT signaling requires its localization with CaN, near sites of Ca^2+^ entry (such as LTCCs), via interactions with A-kinase anchoring protein (AKAP) 79/150 (human 79; rodent 150) (20–22), yet it is unclear if AKAP similarly coordinates CaN-NFAT signaling during iLTP.

Here we show that AKAP-CaN-NFATc3 signaling relays moderate NMDAR activation to the long-term strengthening of inhibitory synapses during iLTP. This represents a novel function for these signaling molecules in promoting protein synthesis and the clustering of receptors and gephyrin to maintain increases in inhibitory synaptic strength. Together, our findings characterize an E-T coupling pathway induced by a moderate NMDA stimulus that controls transcriptional regulation during iLTP stimulation, leading to sustained strengthening of inhibitory synapses.

## RESULTS

### NFATc3 signaling controls sustained inhibitory synaptic potentiation

Our previous work indicated a potential role for NFAT signaling in regulating the expression of inhibitory synaptic proteins during iLTP (6, 7). To test the importance of NFATc3 for iLTP, we used a short hairpin RNA (shRNA) construct to target and knock-down (KD) endogenous NFATc3 (NFATc3 KD) (6, 23) in cultured rat hippocampal neurons and performed electrophysiological recordings of miniature inhibitory postsynaptic currents (mIPSCs). We induced iLTP through a well-characterized chemical paradigm (20 µM NMDA, 10 µM CNQX for 2 min) which rapidly increases inhibitory synapse strength (14–16) and maintains synaptic potentiation out to 90 minutes following induction (6, 7). Recordings in shCtrl (Ctrl) neurons revealed an increase in mIPSC frequency in iLTP-treated neurons compared to sham controls (Fig. 1A-C), but no detectable change in mIPSC amplitude (Fig. 1A,D-E). In contrast to Ctrl neurons, the iLTP-induced increase in mIPSC frequency was prevented in NFATc3 KD neurons (Fig. 1A-C). This effect was rescued upon expression of human NFATc3 (Resc.) which is resistant to the rodent-specific shRNA construct (Fig.1A-C). Notably, NFATc3 KD did not impair mIPSC frequency and amplitude under basal conditions, nor mIPSC kinetics compared to Ctrl or Resc. neurons (Figs. 1A-E, S1A-B), indicating that NFATc3 specifically regulates strengthening of inhibition during iLTP and not basal inhibitory synaptic transmission. In addition, there was no impact of NFATc3 KD or Resc. on basal excitatory synaptic transmission (Fig. S1C-G), suggesting that the effects of NFATc3 loss-of-function are unlikely to be due to alterations to excitatory synaptic proteins. Together, these data indicate a specific requirement for NFATc3 function in the maintenance of iLTP.

**Figure 1.**
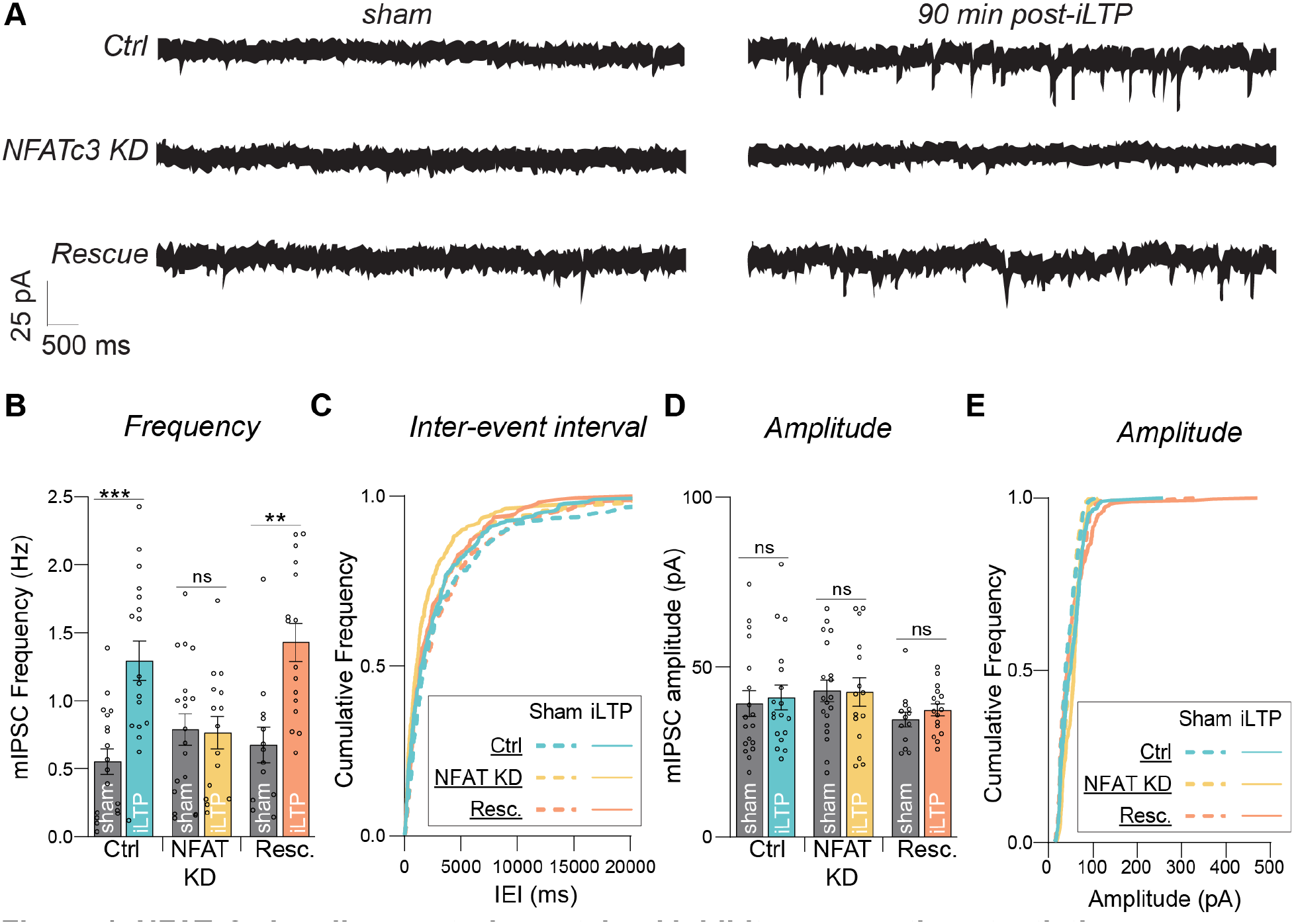
NFATc3 signaling controls sustained inhibitory synaptic potentiation. **(A)** Representative miniature inhibitory postsynaptic current (mIPSC) traces from neurons expressing shCtrl (Ctrl), shNFATc3 (NFATc3 KD), or Rescue (Resc.) in sham or iLTP conditions. **(B)** Quantification of mIPSC frequency from Ctrl, NFAT KD, or Resc. in sham or iLTP conditions. N = 6/n = 12-19 neurons per condition. (main effect of treatment, F(1, 93) = 23.00, p < 0.0001; main effect of condition, F(2, 93) = 2.334, p = 0.1026; interaction, F(2, 93) = 6.426, p = 0.0024; Two-way ANOVA with Tukey’s multiple comparisons post-hoc test). **(C)** Cumulative frequency distribution of mIPSC inter-event intervals (IEIs) from neurons expressing Ctrl, NFAT KD, or Resc. in sham or iLTP conditions. **(D)** Quantification of mIPSC amplitude from Ctrl, NFAT KD, or Resc. in sham or iLTP conditions. N = 6/n = 12-19 neurons per condition. (main effect of treatment, F(1, 93) = 0.2762, p = 0.6005; main effect of condition, F(2, 93) = 1.958, p = 0.1469; interaction, F(2, 93) = 0.1097, p = 0.8962; Two-way ANOVA with Tukey’s multiple comparisons post-hoc test). **(E)** Cumulative frequency distribution of mIPSC amplitude from neurons expressing Ctrl, NFAT KD, or Resc. in sham or iLTP conditions. N = # independent experiments/cultures; n = # neurons. All values represent mean ± SEM. * p < 0.05, *** p < 0.005; 2-way ANOVA with Tukey’s multiple comparisons post-hoc test **(B,D)**.

### NFATc3 is required for increased gephyrin and GABA_A_R clustering following iLTP stimulation

Strengthening of inhibitory synapses during iLTP requires increased clustering of GABA_A_Rs and gephyrin at inhibitory postsynaptic sites (14). If NFATc3 controls the availability of these key synaptic proteins, we would expect NFATc3 KD to impair their accumulation at synapses following iLTP stimulation. To test the requirement of NFATc3 in iLTP-driven gephyrin synaptic clustering, we used live-imaging of cultured neurons expressing a gephyrin intrabody (GFP-Geph IB) to label endogenous gephyrin (24, 25), and live-labeled GABAergic presynaptic terminals with antibodies to vesicular GABA transporter (VGAT), visualized with Oyster^650^. As expected in Ctrl neurons, gephyrin synaptic cluster intensity increased substantially over time following iLTP stimulation compared to sham treatment (Fig. 2A-B). However, NFATc3 KD abolished this increase in gephyrin intensity, compared to Ctrl and Resc. conditions (Fig. 2A-B). Gephyrin cluster area and density were also significantly increased at 90 minutes post-iLTP, an effect that was also prevented by NFATc3 KD, but not in Resc. conditions (Fig. 2C-D).

**Figure 2.**
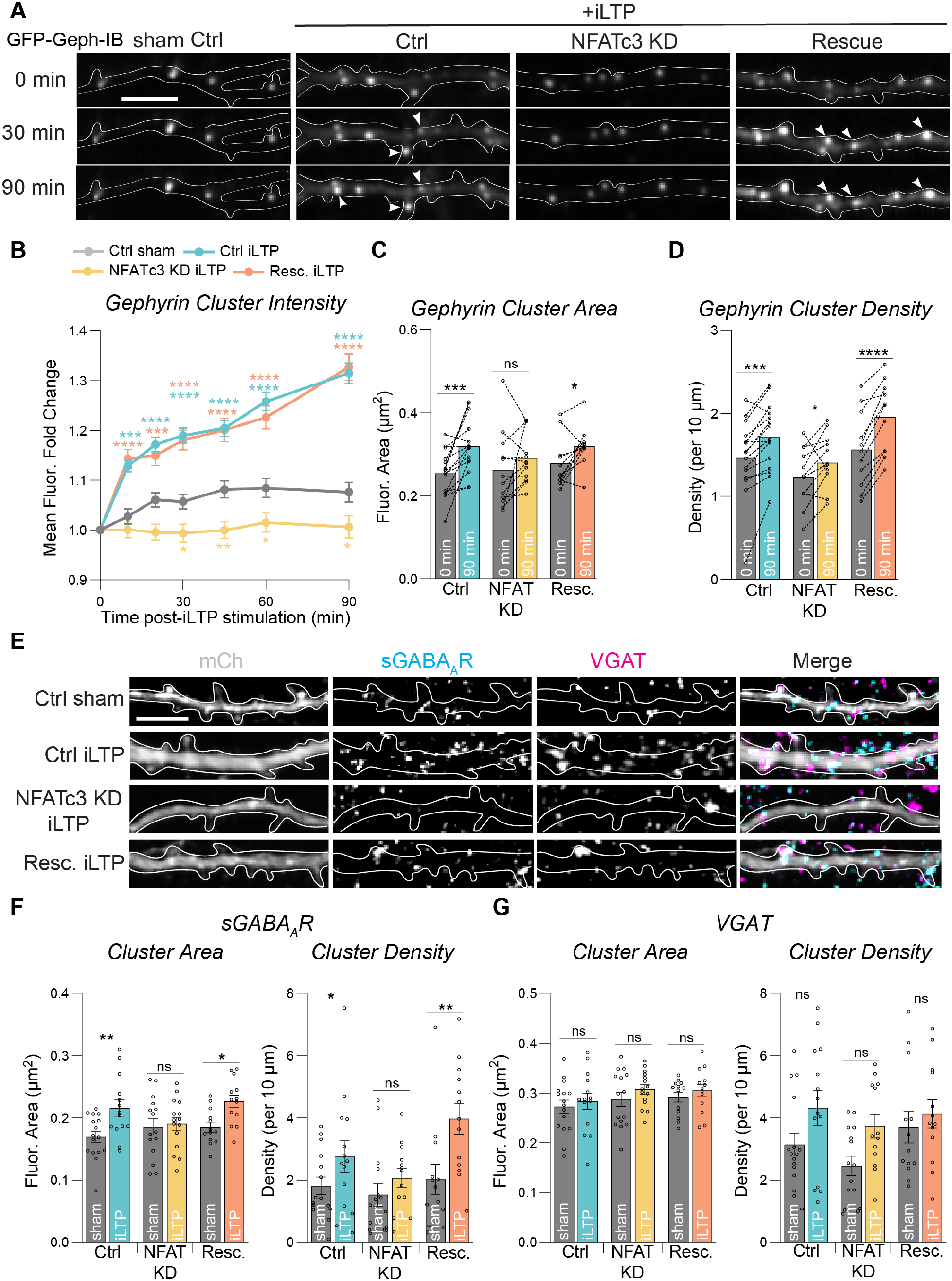
NFATc3 is required for increased gephyrin and GABA_A_R clustering following iLTP stimulation. **(A)** Representative dendritic segment of neurons expressing Ctrl, NFAT KD or Resc. over time in sham or iLTP conditions. Neurons are co-expressing GFP-gephyrin intrabody (GFP-Geph IB). White arrows represent puncta where fluorescence increases over time. Scale bar = 10 µm. **(B)** Quantification of fold change of gephyrin puncta intensity over time following iLTP stimulation in neurons from **(A)**. N = 3/ n = 18 neurons per condition. (Main effect of treatment over time, F(3.655, 2136) = 122.3, p < 0.0001, Geisser-Greenhouse ε = 0.6091; main effect of condition, F(3, 586) = 39.83, p < 0.0001; interaction, F(10.96, 2136) = 23.85, p < 0.0001, Geisser-Greenhouse ε = 0.6091; Mixed-effects model with Geisser-Greenhouse correction). **(C)** Paired measurements of gephyrin cluster area from **(A)** at 0 min and 90 min post-stimulation. N = 3/ n = 11-15 neurons per condition. (main effect of treatment, F(1, 35) = 18.63, p = 0.0001; main effect of condition, F(2, 35) = 0.4543, p = 0.6386; interaction, F(2, 35) = 1.1116, p = 0.3389; 2-way ANOVA with Sidak’s multiple comparisons post-hoc test). **(D)** Paired measurements of gephyrin cluster density from **(A)** at 0 min and 90 min post-iLTP stimulation. N = 3/ n = 11-15 neurons per condition. (main effect of treatment, F(1, 35) = 53.82, p < 0.0001; main effect of condition, F(2, 35) = 3.679, p = 0.0355; interaction, F(2, 35) = 3.065, p = 0.0594; 2-way ANOVA with Sidak’s multiple comparisons post-hoc test). **(E)** Representative dendritic segments of neurons expressing mCh-containing Ctrl, NFAT KD or Resc. in sham or 90 min post-iLTP. Neurons are labeled with antibodies to surface GABA_A_R-γ2 (sGABA_A_R) and VGAT. Scale bar = 5 µm. **(F)** Quantification of sGABA_A_R cluster area (left) and cluster density (right) from **(E)**. N = 4/ n = 12-15 neurons per condition. (<u>Cluster area</u>: main effect of treatment, F(1, 73) = 18.48, p < 0.0001; main effect of condition, F(2, 73) = 1.468, p = 0.2370, interaction, F(2,73) = 3.930, p = 0.0239; <u>Cluster density</u>: main effect of treatment, F(1, 69) = 21.08, p < 0.0001; main effect of condition, F(2, 69) = 4.519, p = 0.0143, interaction, F(2, 69) = 2.423, p = 0.0961; 2-way ANOVA with Tukey’s multiple comparisons post-hoc test). **(G)** Quantification of VGAT cluster area (left) and cluster density (right) from **(E).** N = 4/ n = 12-15 neurons per condition. (<u>Cluster area</u>: main effect of treatment, F(1, 69) = 1.962, p = 0.1651; main effect of condition, F(2, 69) = 1.606, p = 0.2071; interaction, F(2, 69) = 0.06582, p = 0.9364; <u>Cluster density</u>: main effect of treatment, F(1, 69) = 7.473, p = 0.0077; main effect of condition, F(2, 69) = 1.953, p = 0.1488; interaction, F(2, 69) = 0.5539, p = 0.5770; 2-way ANOVA with Tukey’s multiple comparisons post-hoc test). N = # independent experiments/cultures; n = # neurons. All values represent mean ± SEM. * p < 0.05, ** p < 0.01, *** p < 0.005, **** p < 0.0001; Mixed-effects model with Geisser-Greenhouse correction **(B)**; 2-way ANOVA with Sidak’s multiple comparisons post-hoc test **(C-D)**; 2-way ANOVA with Tukey’s multiple comparisons post-hoc test **(F-G)**.

Importantly, NFATc3 KD did not impair pre-treatment cluster area and density of gephyrin compared to Ctrl and Resc. conditions, indicating no impact on basal synaptic clustering (Figs. 2C-D, S2A-C).

Given the impact of NFATc3 KD on synaptic potentiation and increased gephyrin clustering following iLTP, we next tested whether perturbing NFATc3 function would prevent the iLTP-dependent increase in GABA_A_R clustering we had previously observed (7). At 90 minutes following iLTP, neurons were fixed and labeled for surface GABA_A_R-γ2 (sGABA_A_R) and VGAT and imaged with confocal microscopy (Fig. 2E). Cluster analysis revealed increased sGABA_A_R cluster area and density at 90 minutes post-iLTP compared to sham treatments in both Ctrl and Resc. conditions; however, this increase was blocked with NFATc3 KD (Fig. 2E-F). Again, NFATc3 KD did not impact the size and number of sGABA_A_R clusters at inhibitory synapses under basal conditions (Fig. 2E-F), mirroring our results from gephyrin imaging. Analysis of VGAT clusters demonstrated no significant iLTP-induced changes to cluster area and density in any condition (Fig. 2E,G), indicating that NFATc3 KD likely impacts proteins in the postsynaptic domain. Thus, these data demonstrate that NFATc3 is required for the sustained increase in gephyrin, and GABA_A_R clustering following iLTP stimulation and suggest that NFATc3 could control the availability of inhibitory synaptic proteins which are needed to support long-term increases in synapse strength.

### NFATc3 is required for synthesis of synaptic proteins following iLTP stimulation

Late stages of iLTP are characterized by the upregulation of GABA_A_R and gephyrin protein levels which is dependent on increased mRNA translation (7). As NFATc3 has an important role in increased gephyrin and GABA_A_R synaptic clustering following iLTP, we hypothesized that NFATc3 could drive the activity-dependent synthesis of synaptic proteins. To test this hypothesis, we infected hippocampal neurons with Ctrl or NFATc3 KD shRNA lentiviral constructs (23) induced iLTP and measured total levels of inhibitory postsynaptic proteins using western blotting. As we previously showed, following iLTP we observed significantly increased total protein levels for gephyrin and the principal synaptic GABA_A_R subunits α1, γ2, and β3 (Fig. 3A-B) (7). Strikingly, in NFATc3 KD neurons, iLTP-induced increases in gephyrin and GABA_A_R subunit expression were prevented (Fig. 3A-B). Importantly, NFATc3 KD had no impact on gephyrin or GABA_A_R subunit protein expression, nor their total mRNA levels under basal conditions (Fig. 3A-C), indicating that NFATc3 functions specifically during plasticity and does not alter mRNA availability. We also found no change in protein expression of the NMDAR subunit GluN2B following iLTP, nor any basal differences in GluN2B levels with NFATc3 KD compared to Ctrl (Fig. S3A-B), indicating that NMDAR signaling remains intact both during plasticity and in NFATc3 loss-of-function. Taken together, these results suggest that the translation-dependent increase in gephyrin and GABA_A_R protein levels we observe following iLTP stimulation relies on intact NFATc3 signaling.

**Figure 3.**
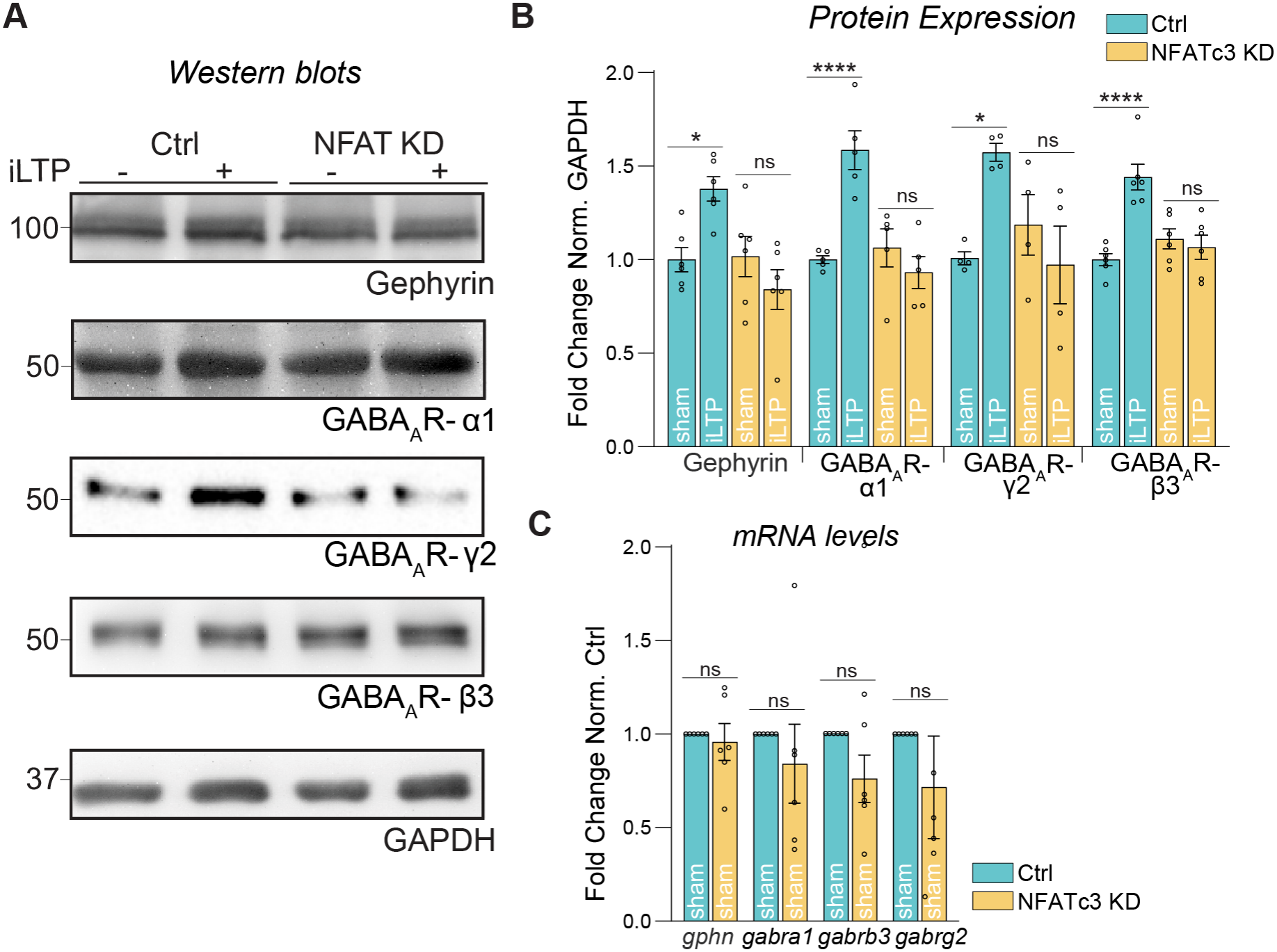
NFATc3 is required for synthesis of inhibitory synaptic proteins following iLTP. **(A)** Western blots showing gephyrin, GABA_A_R subunits α1, γ2, and β3, and GAPDH protein levels in neurons expressing Ctrl (left) or NFATc3 KD (right) either following sham treatment or at 90 minutes post-iLTP. **(B)** Quantification of gephyrin, GABA_A_R subunits α1, γ2, and β3 levels in Ctrl or NFATc3 KD neurons. Protein levels were normalized to GAPDH and data was quantified as relative change in normalized protein expression. n = 4-6 cultures. (Gephyrin: main effect of treatment, F(1, 20) = 1.296, p = 0.2685; main effect of condition, F(1, 20) = 8.636, p = 0.0081, interaction, F(1, 20) = 9.856, p = 0.0052; GABA_A_R-α1: main effect of treatment, F(1, 16) = 7.195, p = 0.0164; main effect of condition, F(1, 16) = 12.20, p = 0.0030; interaction, F(1, 16) = 17.98, p = 0.0006; GABA_A_R-γ2: main effect of treatment, F(1, 12) = 1.711, p = 0.2154; main effect of condition, F(1, 12) = 2.465, p = 0.1424; interaction, F(1, 12) = 8.397, p = 0.0134; GABA_A_R-β3: main effect of treatment, F(1, 20) = 12.45, p = 0.0021; main effect of condition, F(1, 20) = 5.548, p = 0.0288; interaction, F(1, 20) = 18.76, p = 0.0003; 2-way ANOVA with Tukey’s multiple comparisons post-hoc test). **(C)** qRT-PCR of *gphn*, *gabra1*, *gabrg2,* and *gabrb3* mRNA levels in Ctrl or NFATc3 KD neurons. mRNA levels were normalized to GAPDH levels and fold change from Ctrl was quantified for each mRNA. n = 6 cultures. (*gphn* ns, p = 0.6879; *gabra1* ns, p = 0.4861; *gabrg2* ns, p = 0.1176; *gabrb3* ns, p = 0.3461; 1-sample t-test). N = # independent experiments/cultures; n = # neurons. All values represent mean ± SEM. * p < 0.05, ** p < 0.01, *** p < 0.005, **** p < 0.001; 2-way ANOVA with Tukey’s multiple comparisons post-hoc test **(B)**; 1-sample t-test **(C)**.

### NFATc3 enters the nucleus at early timepoints following iLTP induction in a CaN- and LTCC-dependent manner

Following elevated activity levels that lead to high levels of Ca^2+^ influx, NFATc3 rapidly translocates to the nucleus in a CaN-mediated, dephosphorylation-dependent manner and alters gene expression to support excitatory synaptic plasticity (17–19). We previously showed that protein synthesis and increased synaptic clustering of GABA_A_Rs and gephyrin during iLTP were dependent on CaN activity (6, 7), suggesting that a similar CaN-dependent translocation of NFATc3 may also occur during iLTP. To investigate this idea further, we used live-imaging to measure the nuclear abundance of GFP-NFATc3 in hippocampal neurons following iLTP (Fig. 4A). We quantified the change in GFP fluorescence in the nucleus over time following iLTP stimulation and observed NFATc3 enter the nucleus at ∼3 minutes and reside there until ∼13 minutes post-iLTP, with no re-entry at later timepoints (Fig. 4A-B). Application of the CaN inhibitor cyclosporin A (CsA) prevented NFATc3 from entering the nucleus post-iLTP stimulation (Fig. 4A-B), indicating NFATc3 nuclear translocation is indeed driven by CaN activation. We also tested whether LTCC activation was also required for iLTP-driven NFATc3 nuclear translocation, due to their role in propagating Ca^2+^ signals from the dendrites to the soma to drive NFATc3 into the nucleus during eLTP stimulation (18). Application of the LTCC inhibitor nimodipine (Nim) prior to iLTP stimulation prevented NFATc3 nuclear translocation following iLTP (Fig. 4C-D), indicating that LTCC activation is required for this process.

**Figure 4.**
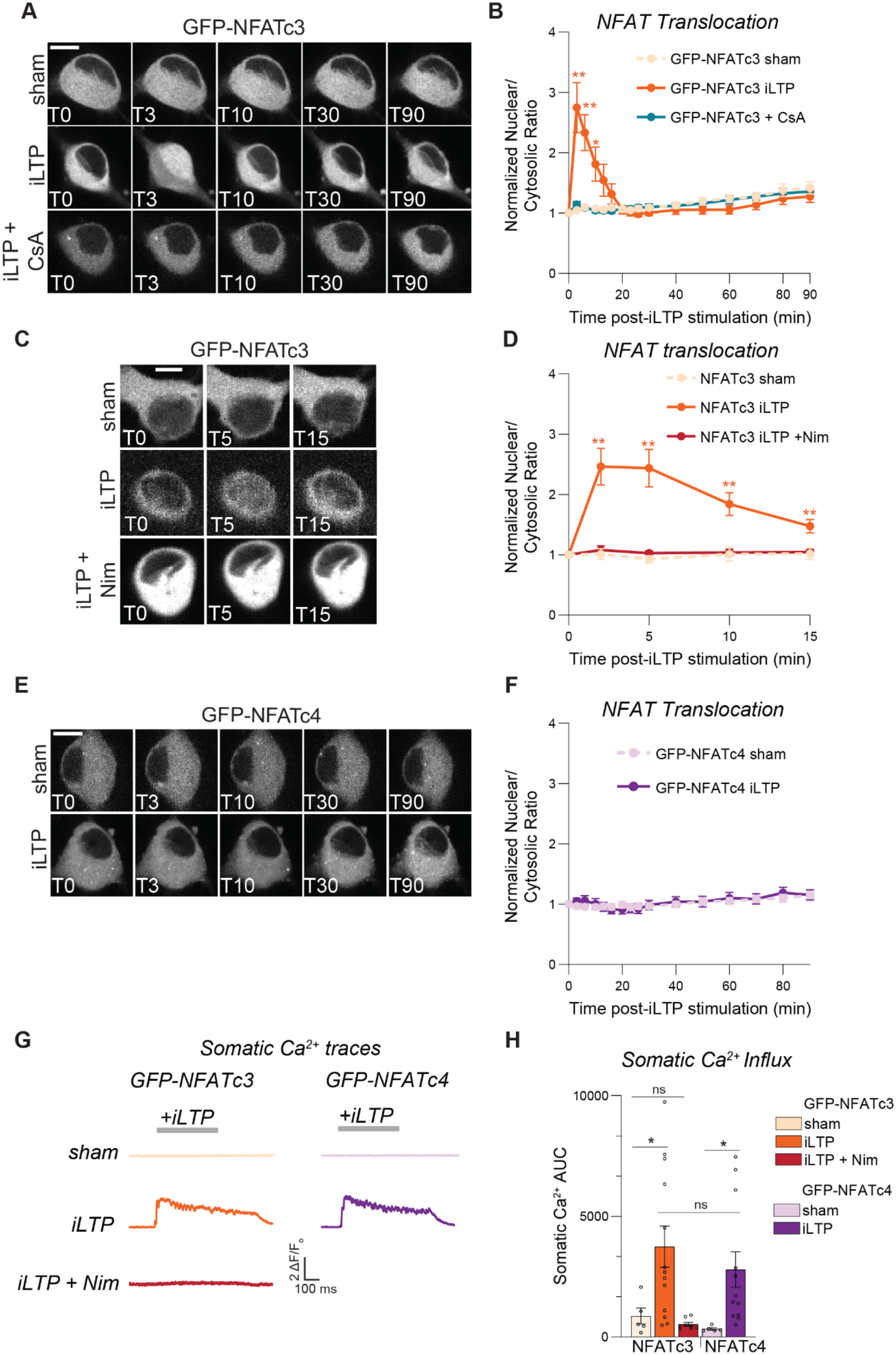
NFATc3 enters the nucleus at early timepoints following iLTP stimulation in a CaN and LTCC-dependent manner. **(A)** Representative images of GFP-NFATc3 translocation over time following sham (top) or iLTP stimulation (middle), with or without the presence of CsA (bottom). Scale bar = 10 µm. **(B)** Quantification of GFP-NFATc3 nuclear to cytosolic intensity normalized to pre-stimulation values over time. N = 3/ n = 10-19 neurons per condition. (main effect of treatment over time, F(2.280, 98.04) = 6.958, p = 0.0009; main effect of condition, F(2, 43) = 3.595, p = 0.0360; interaction, F(4.560, 98.04) = 8.677, p < 0.0001; Repeated measures 2-way ANOVA with the Geisser-Greenhouse correction). **(C)** Representative images of GFP-NFATc3 translocation over time following sham (top) iLTP stimulation with (middle) or without nimodipine (Nim) treatment (bottom). Scale bar = 10 µm. **(D)** Quantification of GFP-NFATc3 nuclear to cytosolic intensity normalized to pre-stimulation values over time. N = 3/ n = 5-13 neurons per condition. (main effect of treatment over time, F(1.465, 33.69) = 5.987, p = 0.0110; main effect of condition, F(2, 23) = 10.76, p = 0.0005; interaction, F(2.929, 33.69) = 7.617, p = 0.0006; Mixed-effects model with Geisser-Greenhouse correction). **(E)** Representative images of GFP-NFATc4 translocation over time following sham (top) or iLTP stimulation (bottom). Scale bar = 10 µm. **(F)** Quantification of GFP-NFATc4 nuclear to cytosolic intensity normalized to pre-stimulation values over time. N = 3/ n = 10-16 neurons per condition. (main effect of treatment over time, F(4.549, 109.2) = 9.111, p = < 0.0001; main effect of condition, F(1, 24) = 0.03698, p = 0.8491; interaction, F(4.459, 109.2) = 1.247, p = 0.2937; Repeated measures 2-way ANOVA with the Geisser-Greenhouse correction). **(G)** Representative Ca^2+^ traces during iLTP stimulation in neurons expressing GFP-NFATc3 or GFP-NFATc4. Scale bar = 10 µm. **(H)** Quantification of GFP-NFATc3 or GFP-NFATc4 maximum nuclear to cytosolic intensity normalized to pre-stimulation values compared to somatic Ca^2+^ area under the curve (AUC) in either sham, iLTP, or iLTP + Nim treatments. N = 3/n = 5-13 neurons per condition. (main effect of treatment, F(2, 39) = 8.645, p = 0.0008; main effect of condition, F(1, 39) = 1.243, p = 0.2717; 2-way ANOVA with Tukey’s post-hoc test). N = # independent experiments/cultures; n = # neurons. All values represent mean ± SEM. * p < 0.05, ** p < 0.01, *** p < 0.005; Repeated measures 2-way ANOVA with the Geisser-Greenhouse correction **(B, D, F)**, 2-way ANOVA with Tukey’s post-hoc test **(H)**.

The other major NFAT isoform in the hippocampus is NFATc4, which is activated by high Ca^2+^ levels generated by strong and prolonged neuronal stimulation (26). Importantly, different levels of Ca^2+^ influx following neuronal stimulation can activate different isoforms of NFAT, leading to differential gene expression (26, 27). To investigate NFAT isoform-specific activation during iLTP, we also imaged GFP-NFATc4. In contrast to NFATc3, NFATc4 did not enter the nucleus following iLTP stimulation (Fig. 4E-F), indicating that NFATc3 is the main isoform activated during iLTP. This result also suggests that the iLTP stimulation causes enough Ca^2+^ influx to activate NFATc3, but is likely not strong enough to activate NFATc4, which requires higher levels of Ca^2+^ influx (26). To verify this hypothesis, we measured changes in somatic Ca^2+^ during iLTP in neurons expressing GFP-NFATc3 or NFATc4 with the Ca^2+^ indicator, jRGECO1a. Following iLTP stimulation, there was a rapid influx of Ca^2+^ into the soma, driving NFATc3 into the nucleus (Fig. 4G-H), an effect that was blocked by the application of Nim, indicating a requirement for LTCC activation. Importantly, the same level of Ca^2+^ influx post-iLTP did not induce NFATc4 nuclear translocation, underlining the specificity of signaling from Ca^2+^ entry to NFATc3 activation during iLTP (Fig. 4G-H). Overall, these results show that NFATc3 is driven into the nucleus at early timepoints following iLTP induction, a process which requires the activation of both LTCCs and CaN, and places NFATc3 in the perfect location to modulate gene expression during plasticity.

### AKAP-CaN signaling drives NFATc3 translocation during iLTP

Much work has shown that effective NFAT signaling to the nucleus requires the coordinated anchoring of both NFAT and CaN near sites of Ca^2+^ influx, such as LTCCs. This anchoring is mediated via interactions with the multifaceted scaffold, AKAP79/150, a key coordinator of CaN activity downstream of Ca^2+^ influx (20–22, 28, 29). Thus, we next asked whether AKAP79/150 could have a novel role in controlling the signaling that drives iLTP. To answer this question, we took advantage of the finding that disrupted binding of CaN to AKAP prevents NFATc3 translocation to the nucleus and utilized previously developed CaN-anchoring deficient AKAP150ΔPIX knock-in mice (ΔPIX), via deletion of the CaN binding PxIxIT motif (30, 31). To initially verify that AKAP-CaN binding acts to drive NFATc3 signaling during iLTP, we imaged cultured WT or ΔPIX mouse hippocampal neurons expressing GFP-NFATc3 and imaged NFATc3 localization at increasing timepoints following iLTP stimulation (Fig. 5A). In WT neurons, NFATc3 was present in the nucleus at 3 and 10 min post-iLTP and localized in the cytosol 20-90 minutes post-iLTP (Fig. 5A-B), confirming similar iLTP-driven translocation in mouse neurons as in rat neurons (Fig. 4A-B). In contrast, in ΔPIX neurons, NFATc3 was not localized to the nucleus at any time following stimulation (Fig. 5A-B), indicating that perturbing CaN binding to AKAP disrupts NFATc3 movement into the nucleus during iLTP.

**Figure 5.**
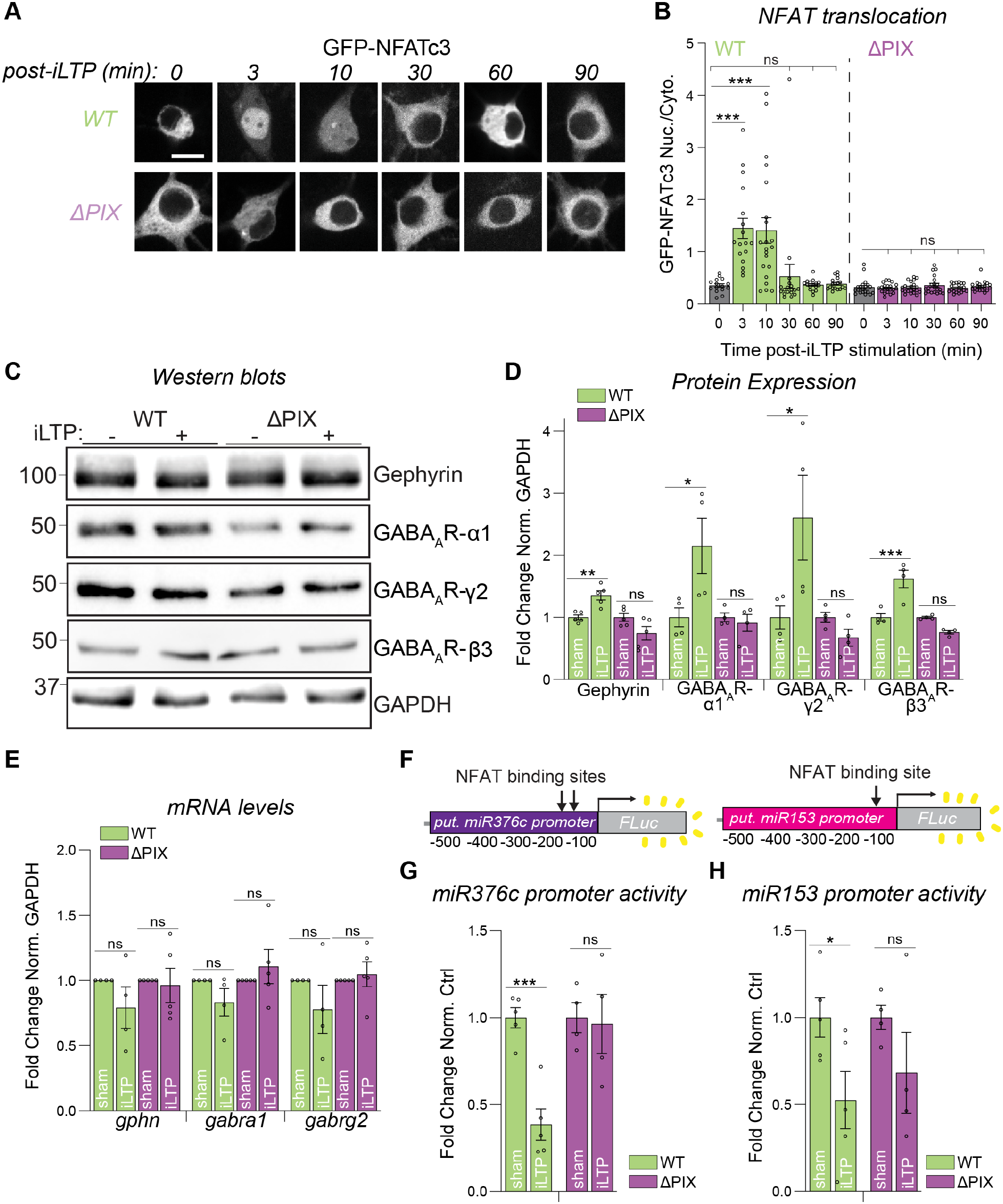
AKAP-CaN binding is required for miRNA transcriptional repression and *de novo* protein synthesis following iLTP. **(A)** Representative images of GFP-NFATc3 translocation in WT or ΔPIX neurons at different timepoints following iLTP induction. Scale bar = 10 µm. **(B)** Quantification of GFP-NFATc3 nuclear to cytosolic intensity at each timepoint following iLTP stimulation in WT and ΔPIX neurons. N = 5/ n = 16-21 neurons per condition. (WT: main effect of treatment over time, F(5, 99) = 9.842, p < 0.0001; ΔPIX: main effect of treatment over time, F(5, 116) = 0.6027, p = 0.6979; 1-way ANOVA with Bonferroni’s post-hoc test). **(C)** Western blots of gephyrin, GABA_A_R subunits α1, γ2, and, β3 and GAPDH levels in WT (left) or ΔPIX (right) neurons either following sham treatment or at 90 minutes post-iLTP. **(D)** Quantification of gephyrin, GABA_A_R subunits α1, γ2, and β3 levels in WT or ΔPIX neurons. Protein levels were normalized to GAPDH, and data was quantified as relative change in normalized protein expression. n = 4-5 cultures. (Gephyrin: main effect of treatment, F(1, 16) = 0.4315; p = 0.5206, main effect of genotype, F(1, 16) = 16.66, p = 0.0009; GABA_A_R-α1: main effect of treatment, F(1, 12) = 4.563; p = 0.0540, main effect of genotype, F(1, 12) = 0.0284, p = 0.0284; GABA_A_R-γ2: main effect of treatment, F(1, 12) = 3.382; p = 0.0908, main effect of genotype, F(1,12) = 7.729, p = 0.0166; GABA_A_R-β3: main effect of treatment, F(1, 12) = 5.783; p = 0.0332, main effect of genotype, F(1, 12) = 29.59, p = 0.0002; 2-way ANOVA with Tukey’s post-hoc test). **(E)** qRT-PCR of *Gphn*, *Gabra1,* and *Gabrg2* mRNA levels in WT and ΔPIX neurons. mRNA levels were normalized to GAPDH levels. n = 4 cultures. (*gphn*: WT sham to WT iLTP ns, p = 0.2823, ΔPIX sham to ΔPIX iLTP ns, p = 0.7926; *gabra1*: WT sham to WT iLTP ns, p = 0.2096, ΔPIX sham to ΔPIX iLTP ns, p = 0.4597; *gabrg2*: WT sham to WT iLTP ns, p = 0.3147, ΔPIX sham to ΔPIX iLTP ns, p = 0.6512; Welch’s t-test). **(F)** Schematic of luciferase constructs for miR376c and miR153 upstream regulatory regions, containing the 500 base pair regulatory region and a Firefly luciferase gene. **(G)** Quantification of miR376c-Luc activity in WT or ΔPIX neurons in sham or 90 min post-iLTP stimulation. Firefly was normalized to Renilla and data was quantified as relative change in normalized activity with error-corrected control values. n = 4 cultures. (WT sham to WT iLTP ***, p = 0.0024; ΔPIX sham to ΔPIX iLTP ns, p = 0.6705; Welch’s t-test). **(H)** Quantification of miR153-Luc activity in WT or ΔPIX neurons in sham or 90 min post-iLTP stimulation. Firefly was normalized to Renilla and data was quantified as relative change in normalized activity with error-corrected control values. n = 4 cultures. (WT sham to WT iLTP *, p = 0.0441; ΔPIX sham to ΔPIX iLTP ns, p = 0.2696; Welch’s t-test). N = # independent experiments/cultures; n = # neurons. All values represent mean ± SEM. ns nonsignificant, * p < 0.05, ** p < 0.01, *** p < 0.005; 1-way ANOVA with Bonferroni’s post-hoc test **(B)**; 2-way ANOVA with Tukey’s post-hoc test **(D);** Welch’s t-test **(E,G-H)**.

### AKAP-CaN binding is required for miRNA transcriptional repression and increased protein expression during iLTP

Given that the AKAP-CaN interaction is required for NFATc3 nuclear translocation and that NFATc3 function is required for the synthesis of gephyrin and GABA_A_R subunits following iLTP, we reasoned that perturbing AKAP-anchored CaN would disrupt the mechanisms driving sustained iLTP. To determine the impact of impaired AKAP-CaN signaling on inhibitory protein synthesis, we used western blotting in either WT or ΔPIX hippocampal neurons to measure total protein levels. In WT neurons we observed an increase in gephyrin, GABA_A_R-α1, GABA_A_R-γ2, and GABA_A_R-β3 protein levels 90 minutes following iLTP compared to sham treatments (Fig. 5C-D). However, this iLTP-induced increase in protein expression was not observed in ΔPIX neurons (Fig. 5C-D). Perturbing AKAP-CaN binding also did not impact inhibitory postsynaptic protein levels under basal conditions (Fig. 5C-D), indicating that this interaction is specifically required for the iLTP-dependent increase in protein expression. Furthermore, qRT-PCR demonstrated no change in *gphn*, *gabra1*, or *gabrg2* mRNA levels in WT or ΔPIX neurons under basal conditions or following iLTP (Fig. 5E), indicating that the changes we observed to total protein levels are likely due to increased translation.

Reduced transcription of miR376c and miR153 is required for the translation of gephyrin and GABA_A_R subunit mRNAs to increase total protein levels following iLTP (6, 7). We previously identified active NFATc3 binding sites in upstream regulatory regions of these miRNAs that are required to reduce their transcriptional activity following iLTP stimulation (6, 7). To test the requirement of AKAP-CaN binding for this transcriptional repression, we used reporter plasmids where Firefly luciferase (Luc) is fused to either the miR376c or miR153 regulatory region, in which Luc expression acts as a readout of miRNA promoter activity (6, 7) (Fig. 5F). In WT neurons, at 90 minutes post-iLTP there was a robust reduction in Luc luminescence indicating a reduction in miR376c and miR153 transcription (Fig. 5G-H), in agreement with our previous work (6, 7). However, no change in regulatory region activity was observed for either miRNA in ΔPIX neurons (Fig. 5G-H), indicating that AKAP-CaN signaling is likely pivotal in controlling the iLTP-induced transcriptional repression of miR376c and miR153. Together, these data show that AKAP-anchored CaN engages NFATc3 signaling within this miRNA-mediated pathway to increase inhibitory synaptic protein production following iLTP stimulation.

### AKAP-anchored CaN is required for increased gephyrin and GABA_A_R clustering following iLTP

Given the importance of the AKAP-CaN interaction for the iLTP-induced synthesis of inhibitory synaptic proteins, we next asked if AKAP-CaN signaling also drives the long-term increase of gephyrin and GABA_A_R clustering following iLTP. To address this question, we evaluated gephyrin and GABA_A_R synaptic clustering in WT and ΔPIX mouse hippocampal neurons, labeling for gephyrin, sGABA_A_R, and VGAT followed by confocal imaging (Fig. 6A). As expected, cluster analysis revealed that at 90 minutes post-iLTP, WT neurons exhibited increased gephyrin and sGABA_A_R cluster area and density compared with sham (Fig. 6A-B,E-F). In contrast, in ΔPIX neurons, iLTP-induced changes in gephyrin and sGABA_A_R clustering were prevented (Fig. 6A-C,E-F). There were no changes in VGAT cluster area or density 90 minutes following iLTP in WT or ΔPIX neurons (Fig. 6A,D,G). Moreover, we observed no difference in gephyrin, sGABA_A_R, or VGAT clustering in either WT or ΔPIX neurons under basal conditions (Fig. 6A-G), suggesting that perturbation of AKAP-CaN binding only impacts synaptic clustering following iLTP.

**Figure 6.**
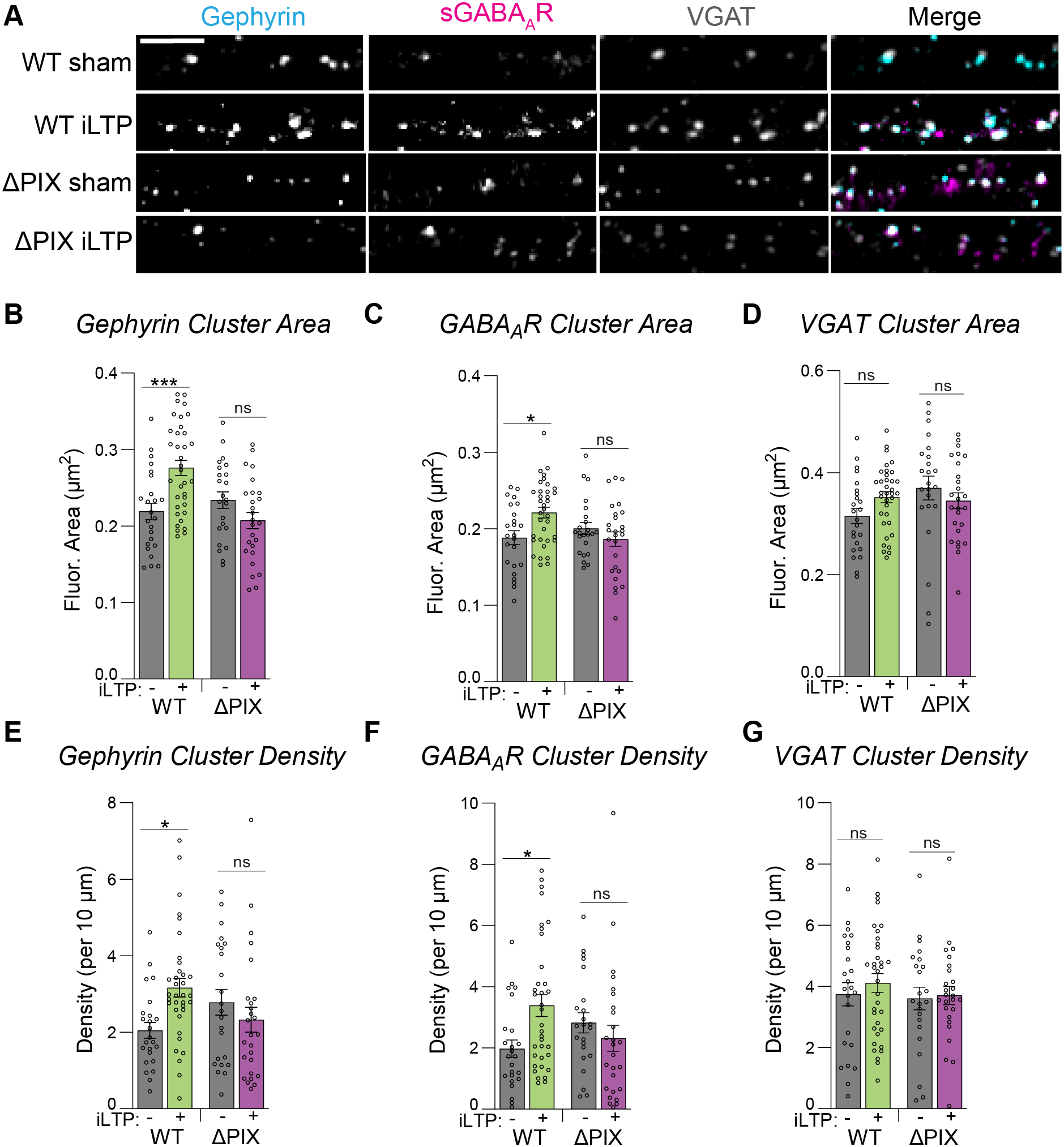
AKAP-anchored CaN is required for increased gephyrin and GABA_A_R clustering following iLTP. **(A)** Representative dendritic segments of WT and ΔPIX neurons following sham treatment or at 90 min post-iLTP. Neurons labeled with antibodies to gephyrin, surface GABA_A_R-γ2 (sGABA_A_R), and VGAT. Scale bar = 5 µm. **(B)** Quantification of gephyrin cluster area from **(A)**. N = 3/ n = 23-35 neurons per condition. (main effect of treatment, F(1, 104) = 1.993, p = 0.1610; main effect of condition, F(1, 104) = 6.321, p = 0.0135; interaction, F(1, 104) = 15.35, p = 0.0002; 2-way ANOVA with Tukey’s post hoc test). **(C)** Quantification of sGABA_A_R cluster area from **(A)**. N = 3/ n = 23-35 neurons per condition. (main effect of treatment, F(1, 104) = 1.266, p = 0.2632; main effect of condition, F(1, 104) = 1.816, p = 0.1807; interaction, F(1, 104) = 7.802, p = 0.0062; 2-way ANOVA with Tukey’s post hoc test). **(D)** Quantification of VGAT cluster area from **(A)**. N = 3/ n = 23-35 neurons per condition. (main effect of treatment, F(1, 104) = 0.1397, p = 0.7093; main effect of condition, F(1, 104) = 2.82, p = 0.1339, interaction, F(1, 104) = 3.770, p = 0.0549; 2-way ANOVA with Tukey’s post hoc test). **(E)** Quantification of gephyrin cluster density from **(A)**. N = 3/ n = 23-35 neurons per condition. (main effect of treatment, F(1, 104) = 1.364, p = 0.2455; main effect of condition, F(1, 104) = 0.03498, p = 0.8520; interaction, F(1, 104) = 7.554, p = 0.0071; 2-way ANOVA with Tukey’s post hoc test). **(F)** Quantification of sGABA_A_R cluster density from **(A)**. N = 3/ n = 23-35 neurons per condition. (main effect of treatment, F(1, 104) = 1.530, p = 0.2189; main effect of condition, F(1, 104) = 0.08824, p = 0.7670; interaction, F(1, 104) = 6.787, p = 0.0105; 2-way ANOVA with Tukey’s post hoc test). **(G)** Quantification of VGAT cluster density from **(A)**. N = 3/ n = 23-35 neurons per condition. (main effect of treatment, F(1, 104) = 0.4680, p = 0.4954; main effect of condition, F(1, 104) = 0.6325, p = 0.4282; interaction, F(1, 104) = 0.1537, p = 0.6958; 2-way ANOVA with Tukey’s post hoc test) N = # independent experiments/cultures; n = # neurons. All values represent mean ± SEM. * p < 0.05, *** p < 0.005; 2-way ANOVA with Tukey’s post hoc test **(B-G)**.

Given that one potential caveat of using the ΔPIX mouse model is the possibility of long-term effects of perturbed AKAP-CaN binding and compensation, we complemented these experiments using an acute KD/rescue model. We performed similar confocal imaging experiments as described above but used cultured rat hippocampal neurons transiently transfected with shRNA to KD AKAP150 (AKAP150 KD) and either a human mCh-AKAP79WT (AKAP79WT) or mCh-AKAP79ΔPIX (AKAP79ΔPIX) rescue constructs (Fig. S4A). Consistent with our mouse culture experiments, iLTP-induced increases in gephyrin cluster area and density were blocked in AKAP79ΔPIX expressing rat neurons, with no impact of AKAP79ΔPIX on gephyrin and VGAT clustering under basal conditions (Fig. S4A-E). Thus, these results in rat neurons, along with our experiments in ΔPIX mouse neurons, suggest that AKAP-CaN signaling is critical for the growth and formation of inhibitory synapses during iLTP.

### Inhibitory synaptic potentiation during iLTP requires AKAP-CaN signaling

Finally, we wanted to test whether AKAP-CaN signaling drives the persistent strengthening of synaptic inhibition during iLTP. To assess the functional impact of the ΔPIX mutation on iLTP, we measured mIPSCs in either cultured WT or AKAP150ΔPIX mouse hippocampal neurons at 90 minutes post-iLTP or sham conditions (Fig. 7A). In WT neurons, mIPSC frequency increased during iLTP compared to sham treatments (Fig. 7A-C), with no change in amplitude (Fig. 7A,D-E), similar to our observations in rat neurons (Fig. 1). In contrast, in ΔPIX neurons there was no change in mIPSC frequency following iLTP (Fig. 7A-C), indicating that AKAP-CaN signaling is required for iLTP. Crucially, there were no differences in mIPSC frequency, amplitude, or rise-time between WT and ΔPIX neurons under basal conditions (Fig. 7A-E, S5A). However, there was a significant increase in mIPSC decay time following iLTP in ΔPIX neurons (Fig. S5B), indicating possible differences in GABA_A_R subunit composition post-iLTP (32), which could reflect compensation in these neurons. Altogether, these function data demonstrate that perturbing AKAP-CaN signaling is sufficient to block long-term inhibitory synaptic plasticity, mirroring our mIPSC recordings upon NFATc3 KD and suggesting that AKAP-CaN-NFATc3 signaling controls sustained iLTP.

**Figure 7.**
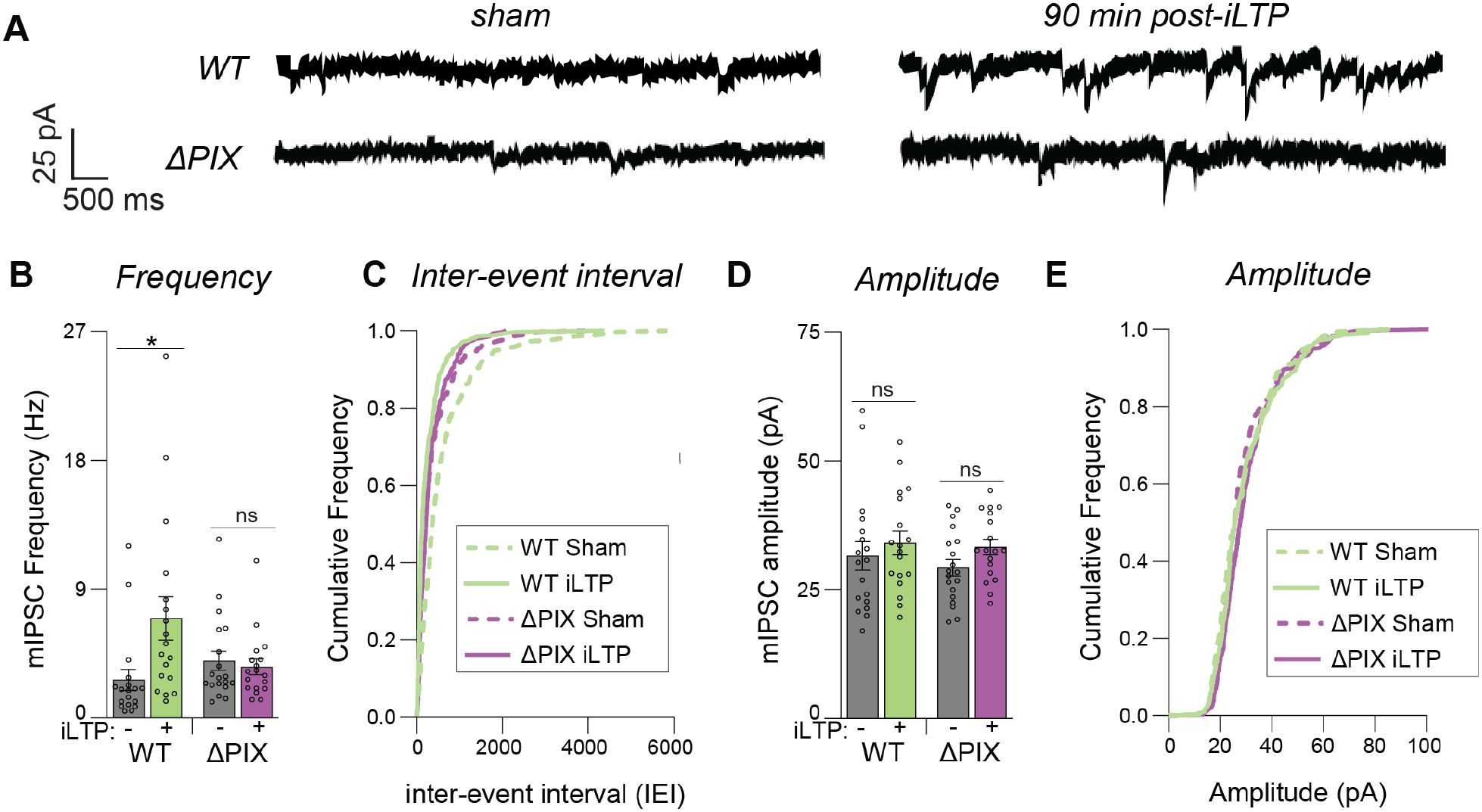
Inhibitory synaptic potentiation during iLTP requires AKAP-CaN signaling. **(A)** Representative miniature inhibitory postsynaptic current (mIPSC) traces from WT and ΔPIX mouse neurons in sham or iLTP conditions. **(B)** Quantification of mIPSC frequency from WT and ΔPIX neurons in sham or iLTP conditions. N = 5/ n =18-19 neurons per condition. (main effect of treatment, F(1, 69) = 4.263, p = 0.0427; main effect of condition, F(1, 69) = 1.207, p = 0.2757; interaction, F(1, 69) = 6.403, p = 0.0137; 2-way ANOVA with Tukey’s post-hoc test). **(C)** Cumulative frequency distribution of mIPSC inter-event intervals (IEIs) from WT and ΔPIX neurons in sham or iLTP conditions. **(D)** Quantification of mIPSC amplitude from WT and ΔPIX neurons in sham or iLTP conditions. N = 5/ n =18-19 neurons per condition. (main effect of treatment, F(1, 69) = 2.382, p = 0.1273; main effect of condition, F(1, 69) = 0.5363, p = 0.4664; interaction, F(1, 69) = 0.1254, p = 0.7244; 2-way ANOVA with Tukey’s post-hoc test). **(E)** Cumulative frequency distribution of mIPSC amplitude from WT and ΔPIX neurons in sham or iLTP conditions. N = # independent experiments/cultures; n = # neurons. All values represent mean ± SEM. * p < 0.05; 2-way ANOVA with Tukey’s post-hoc test **(B,D)**.

## DISCUSSION

Long-term synaptic plasticity requires coordinated changes in gene expression to produce persistent alterations in synaptic strength (10). Decades of work have interrogated the E-T coupling mechanisms that allow neurons to increase excitatory synaptic strength in response to high activity neuronal stimuli (33–35). Yet, how neuronal activity signals alterations in gene expression to control inhibitory synaptic plasticity has remained poorly understood. In our previous work, we demonstrated that transcriptional regulation is also required for sustained increases in inhibitory synapse strength following moderate NMDAR stimulation (6, 7). These findings suggested the existence of an E-T coupling mechanism that is activated in response to a moderate NMDAR stimulation and alters gene expression to support sustained iLTP. Our results support a model (Fig. 8), in which iLTP induction triggers Ca^2+^ influx through LTCCs, activating AKAP-anchored CaN to dephosphorylate NFATc3, permitting its rapid translocation into the nucleus. NFATc3 subsequently contributes to the transcriptional regulation of miR376c and miR153, relieving miRNA-mediated translational repression of GABA_A_R and gephyrin mRNAs and enabling their increased synthesis. The resulting new receptors and scaffolds then cluster at new and existing synapses thereby maintaining the potentiation of synaptic strength. E-T coupling mechanisms have been shown to regulate synaptic inhibition induced by high-activity in behavioral enrichment paradigms (36–38). These mechanisms require high levels of activity, over longer time periods and engage NPAS4 and BDNF which increase synapse number over hours-to-days. In comparison, we provide evidence for a more rapid E-T coupling mechanism that enhances translation of key inhibitory targets, enabling the continued postsynaptic strengthening of inhibition following an acute, more moderate elevation in activity.

**Figure 8.**
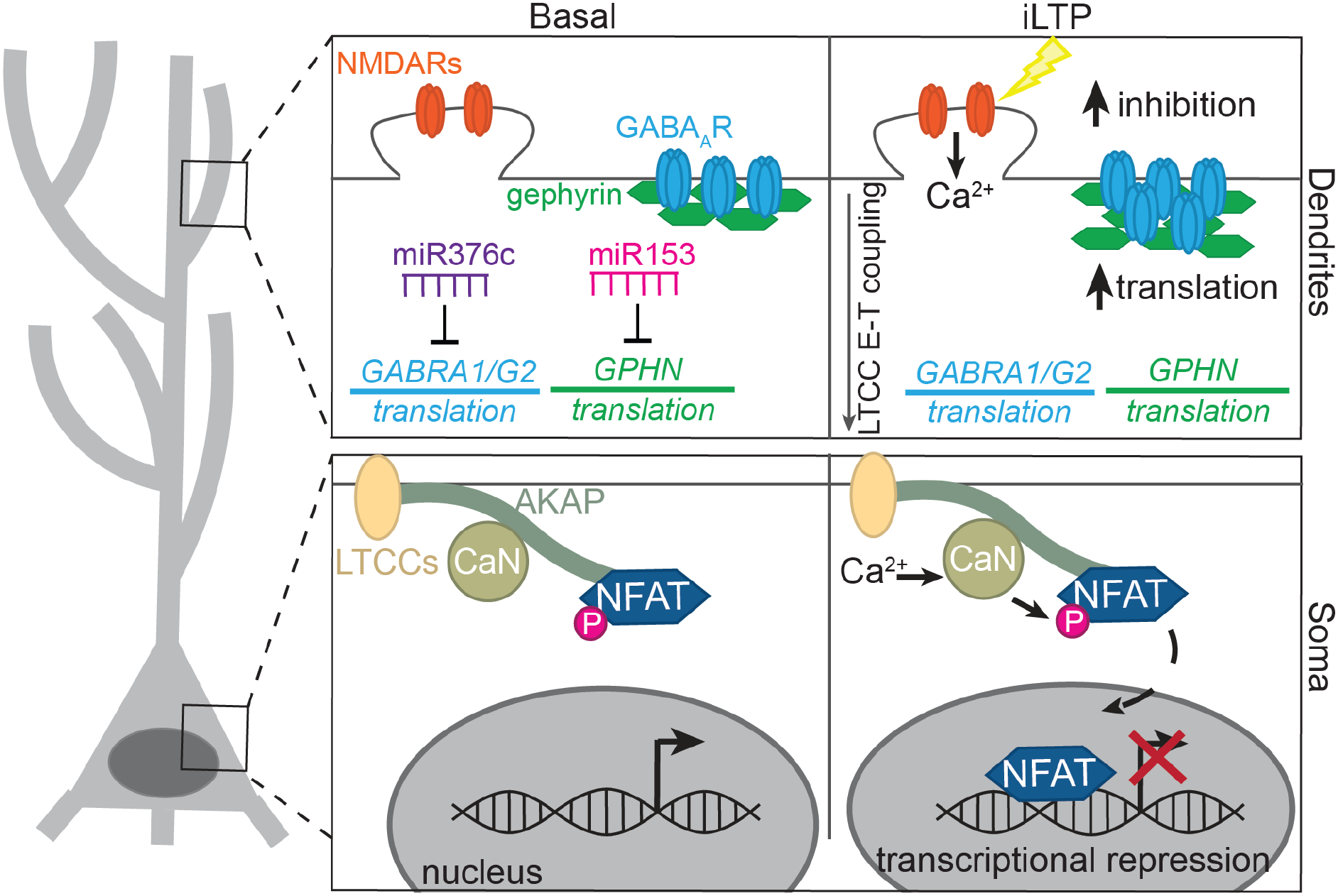
AKAP-CaN-NFATc3 signaling is required for *de novo* protein synthesis for sustained structural and functional iLTP. Schematic of NMDAR-mediated iLTP stimulation inducing NFATc3 translocation, regulated by the AKAP-CaN complex, leading to gene repression and *de novo* synthesis of inhibitory synaptic proteins to sustain persistent iLTP.

We previously identified that transcriptional repression of miR376c and miR153 is required for *de novo* synthesis of gephyrin and GABA_A_Rs during iLTP, thereby supporting long-lasting potentiation of inhibitory synaptic strength (6, 7). We identified active NFAT-binding sites within the upstream regulatory region of miR376c and miR153 genes (6, 7), raising the possibility that NFAT-mediated E-T coupling could regulate iLTP-associated transcriptional changes and upregulation of inhibitory synaptic proteins. Yet whether NFATc3 or its upstream signaling pathway was required for iLTP had not been established. NFATs regulate genes that function in synaptic plasticity, Ca^2+^ regulation, neuronal excitability, and development (21, 39–43). Thus, to dissect the acute, post-developmental and postsynaptic contributions of NFATc3 to iLTP, we used transient and sparse knock-down of NFATc3 in hippocampal neurons. NFATc3 KD prevented the robust increase in mIPSC frequency we observed in Ctrl neurons at 90 minutes post-iLTP, indicating a requirement for NFATc3 in the formation of new inhibitory synapses during plasticity. In addition, our live imaging experiments indicate that following iLTP, gephyrin and GABA_A_R puncta density not only increased in dendrites, but puncta area was also increased, effects that were blocked by NFATc3 KD. This apparent discrepancy between our electrophysiological and imaging results is likely due to iLTP predominantly potentiating dendritic synapses rather than somatic synapses (6, 7, 12). Thus, increased frequency likely reflects the growth of new and existing dendritic synapses (as shown by imaging), but any increases in mIPSC amplitude are masked due to filtering of dendritic mIPSCs and unchanged mIPSC somatic responses (6, 7). Importantly, NFATc3 KD had no impact on inhibitory or excitatory synaptic transmission in sham conditions, suggesting that NFATc3 signaling does not play a significant role in maintaining synaptic transmission under basal conditions.

We previously showed that *de novo* synthesis of GABA_A_R subunits and gephyrin and their subsequent accumulation at GABAergic synapses is required for sustained increases in inhibitory synaptic strength following iLTP induction (7). Here we show that this iLTP-driven *de novo* synthesis requires NFATc3 signaling, suggesting that the mechanism by which NFATc3 contributes to iLTP is via the modulation of mRNA translation. Notably, we also observed an iLTP-induced increase in the expression levels of the GABA_A_R-β3 subunit, a common receptor subunit that forms with α1-and γ2-subunits to form functional receptors. β3-containing

GABA_A_Rs are also essential for the first stages of iLTP that involve rapid trafficking to the synapse (14), thus it follows that more β3 would need to be produced to support β3-containing receptors over longer time periods. NFATc3 KD also prevented the upregulation of β3 total protein levels post-iLTP, suggesting a conserved mechanism by which NFATc3 coordinates the regulation of other inhibitory synaptic proteins for sustained potentiation. Future work is required to dissect the specific mechanisms controlling GABA_A_R-β3 upregulation, as well as determining whether NFATc3 plays a role in the upregulation of other proteins during iLTP.

What mechanisms might direct NFATc3 to specifically induce iLTP-dependent gene regulation? Firstly, our data suggest that iLTP stimulation induces Ca^2+^ transients that are sufficient to activate NFATc3, but not NFATc4, therefore activating a specific set of NFATc3-targeted genes. This selectivity of activation is due to these isoforms exhibiting differential sensitivity to Ca^2+^, CaN-binding affinities, and dephosphorylation efficiencies (26, 27). Secondly, NFATs regulate gene expression in collaboration with other activity-dependent transcription factors, such as the AP-1 complex and STAT3 (40, 44), so it is possible that although NFATc3 is necessary for iLTP, other co-factors may also be required and confer specificity to NFATc3 signaling in the context of iLTP. Finally, we show that NFATc3 represses transcription following iLTP induction (7), which is contrary to the canonical role of NFATs in promoting the transcription of its targets (40). This atypical role for NFATs has been observed in bone cells, immune cells and early in neuronal development (45–47) and it is thought that NFATs induce transcriptional-repression via recruitment of histone deacetylases (HDACs) to specific DNA regions to induce gene silencing (48). Thus, it is tempting to speculate that a similar mechanism could be employed by NFATc3 to specifically represses transcription during iLTP.

Our data identify AKAP-anchored CaN as an upstream regulator of NFATc3 during iLTP. Perturbation of AKAP-CaN interactions or NFATc3 function had almost identical effects on iLTP-induced synaptic growth and potentiation as well as protein synthesis. Importantly, disrupted AKAP-CaN signaling prevented NFATc3 nuclear translocation. Together, these results place AKAP-CaN signaling upstream of NFATc3 activation during iLTP and establish a previously unrecognized role for this signaling complex in inhibitory synaptic plasticity. The involvement of AKAP-anchored CaN in the late-phase of iLTP is of particular interest because AKAP-CaN signaling is also crucial for AMPA receptor (AMPAR) endocytosis during eLTD, a process also induced by moderate NMDAR stimulation (30). Thus, the same NMDAR stimulation may engage AKAP-CaN signaling to produce temporally distinct changes in excitatory and inhibitory synaptic transmission: rapidly weakening excitatory synaptic strength while promoting long-term increases in synaptic inhibition. Such coordinated regulation could provide an effective mechanism to alter dendritic excitability during specific patterns of activity. Previous studies have suggested that AKAP may also play a role directly at inhibitory synapses (49), potentially via direct interaction with the GABA_A_R-β3 subunit (50). Yet through all our experiments we find no significant effect of impaired AKAP-CaN signaling on any measured parameters. Regardless of any possible role for AKAP-CaN at the inhibitory synapse itself, our data establishes that the AKAP-CaN association contributes to iLTP over longer timescales and this is mimicked by NFATc3 KD, which is downstream of AKAP-CaN signaling.

In summary, our findings identify a novel role for AKAP-CaN-NFATc3 signaling in the mechanistic underpinnings of long-term inhibitory postsynaptic plasticity. This form of plasticity is critical for controlling pyramidal cell output and the ability of these cells to undergo eLTP (51–53), as well as hippocampal place cell stability and flexibility (51). Therefore, deciphering molecular mechanisms that control this form of plasticity informs our understanding of cellular mechanisms underlying learning and cognition. Future studies will determine the transcriptional co-factors recruited by NFATc3, establish how NFATc3 mediates repression of miRNAs, and determine how this pathway interacts with other forms of excitatory and inhibitory synaptic plasticity to coordinate neuronal excitability.

---

**Figure S1.**
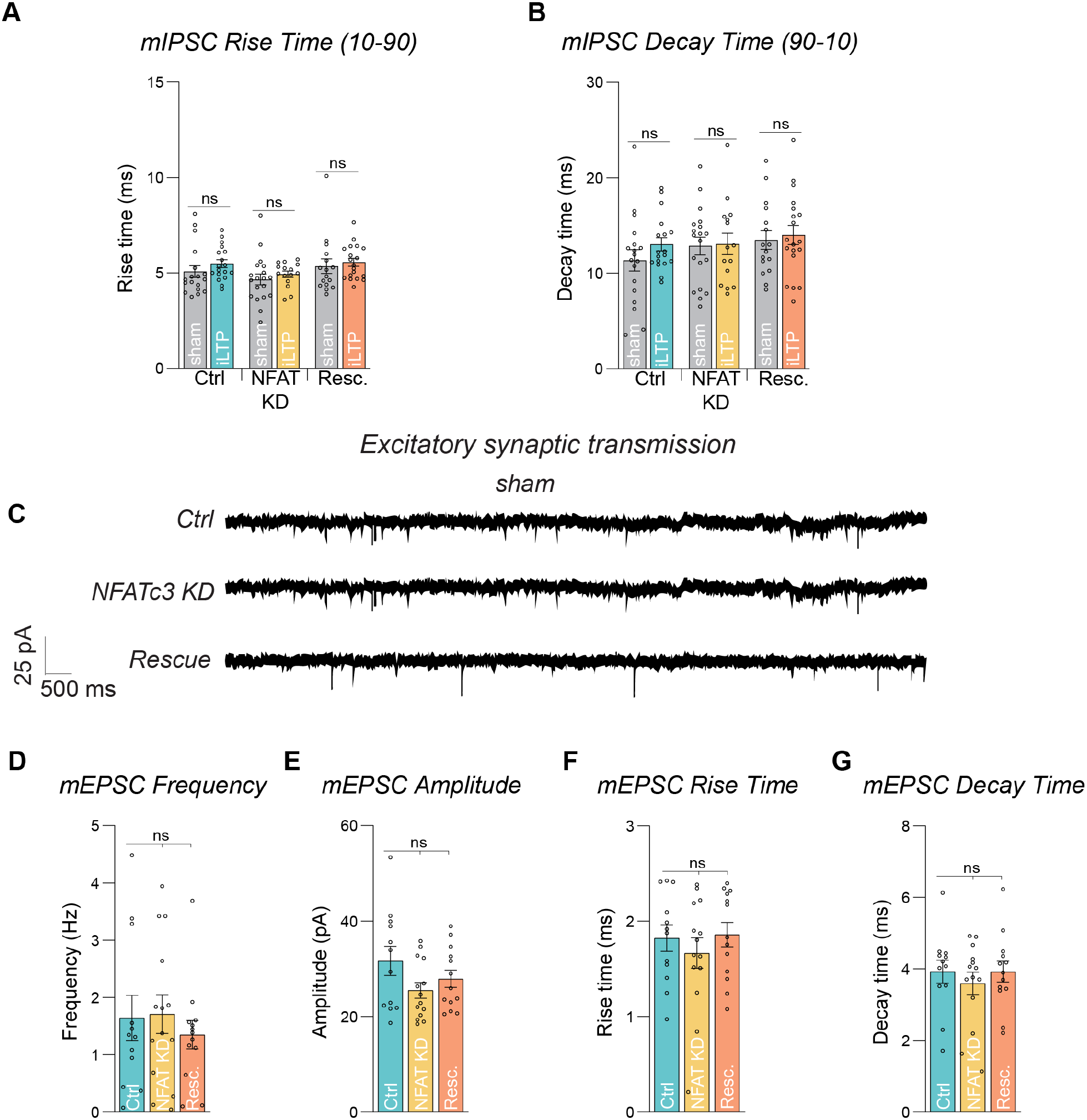
Supplement to Figure 1. NFATc3 KD did not impair mIPSC kinetics or basal excitatory synaptic transmission. **(A)** Quantification of mIPSC rise time from Ctrl, NFAT KD, or Resc. in sham or iLTP conditions. N = 6/n = 15-19 neurons per condition. (main effect of treatment, F(1, 99) = 1.859, p = 0.1759; main effect of condition, F(2, 99) = 3.170, p = 0.0463; interaction, F(2, 99) = 0.08631, p = 0.9174; Two-way ANOVA with Tukey’s multiple comparisons post-hoc test). **(B)** Quantification of mIPSC decay time from Ctrl, NFAT KD, or Resc. in sham or iLTP conditions. N = 6/n = 15-19 neurons per condition. (main effect of treatment, F(1, 99) = 1.017, p = 0.3158; main effect of condition, F(2, 99) = 1.276, p = 0.2836; interaction, F(2, 99) = 0.3187, p = 0.7278; Two-way ANOVA with Tukey’s multiple comparisons post-hoc test). **(C)** Representative miniature excitatory postsynaptic current (mEPSC) traces from neurons expressing Ctrl, NFAT KD, or Resc. in sham conditions. **(D)** Quantification of mEPSC frequency from Ctrl, NFAT KD, or Resc. in sham conditions. N = 5/n = 12-14 neurons per condition. (main effect of treatment, F(2, 36) = 0.3411, p = 0.7133; 1-way ANOVA with Bonferroni’s post-hoc test). **(E)** Quantification of mEPSC amplitude from Ctrl, NFAT KD, or Resc. in sham conditions. N = 5/n = 12-14 neurons per condition. (main effect of treatment, F(2, 36) = 1.991, p = 0.1513; 1-way ANOVA with Bonferroni’s post-hoc test). **(F)** Quantification of mEPSC rise time from Ctrl, NFAT KD, or Resc. in sham conditions. N = 5/n = 12-14 neurons per condition. (main effect of treatment, F(2, 36) = 0.5282, p = 0.5942; 1-way ANOVA with Bonferroni’s post-hoc test). **(G)** Quantification of mEPSC decay time from Ctrl, NFAT KD, or Resc. in sham conditions. N = 5/n = 12-14 neurons per condition. (main effect of treatment, F(2, 36) = 0.3873, p = 0.6817; 1-way ANOVA with Bonferroni’s post-hoc test). N = # independent experiments/cultures; n = # neurons. All values represent mean ± SEM. ns = not significant; 2-way ANOVA with Tukey’s multiple comparisons post-hoc test **(A-B)**; 1-way ANOVA with Bonferroni’s post-hoc test **(D-G)**.

**Figure S2.**
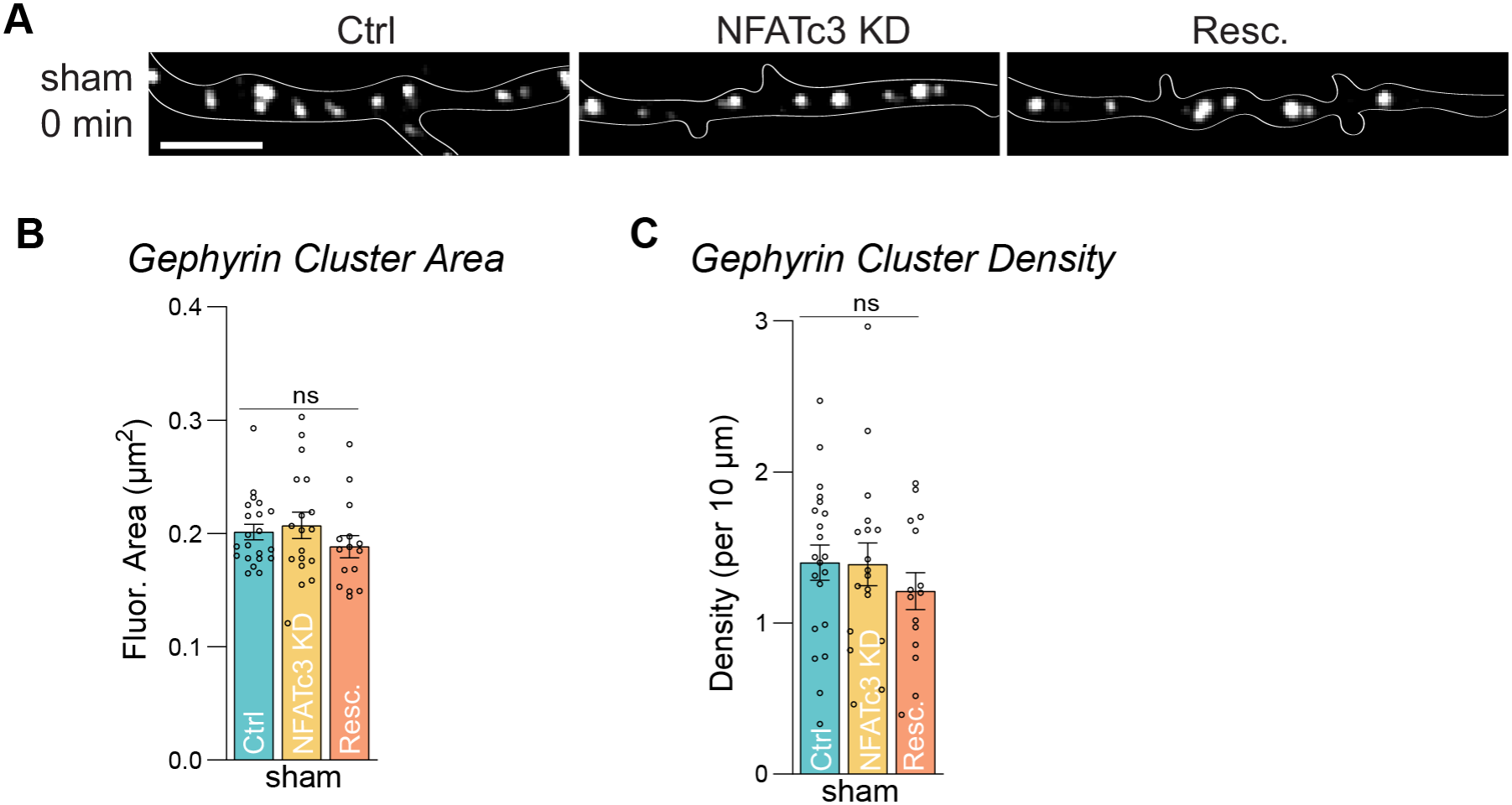
Supplement to Figure 2. NFATc3 KD did not impact basal gephyrin cluster area or cluster density. **(A** Representative dendritic segment of neurons expressing Ctrl, NFAT KD or Resc. over time in sham conditions. Neurons are co-expressing GFP-gephyrin intrabody (GFP-Geph IB). Scale bar = 10 µm. **(B)** Gephyrin cluster area prior to iLTP stimulation cluster area at 0 minutes following sham treatment. N = 3/ n = 15-21 neurons per condition. (main effect of treatment, F(2, 51) = 0.9449, p = 0.3954; One-way ANOVA with Bonferroni’s post-hoc test). **(C)** Gephyrin cluster density prior to iLTP stimulation cluster area at 0 minutes following sham treatment. N = 3/ n = 15-21 neurons per condition. (main effect of treatment, F(2, 51) = 0.6238, p = 0.5399; One-way ANOVA with Bonferroni’s post-hoc test). N = # independent experiments/cultures; n = # neurons. All values represent mean ± SEM. 1-way ANOVA with Bonferroni’s post-hoc test **(B-C)**

**Figure S3.**
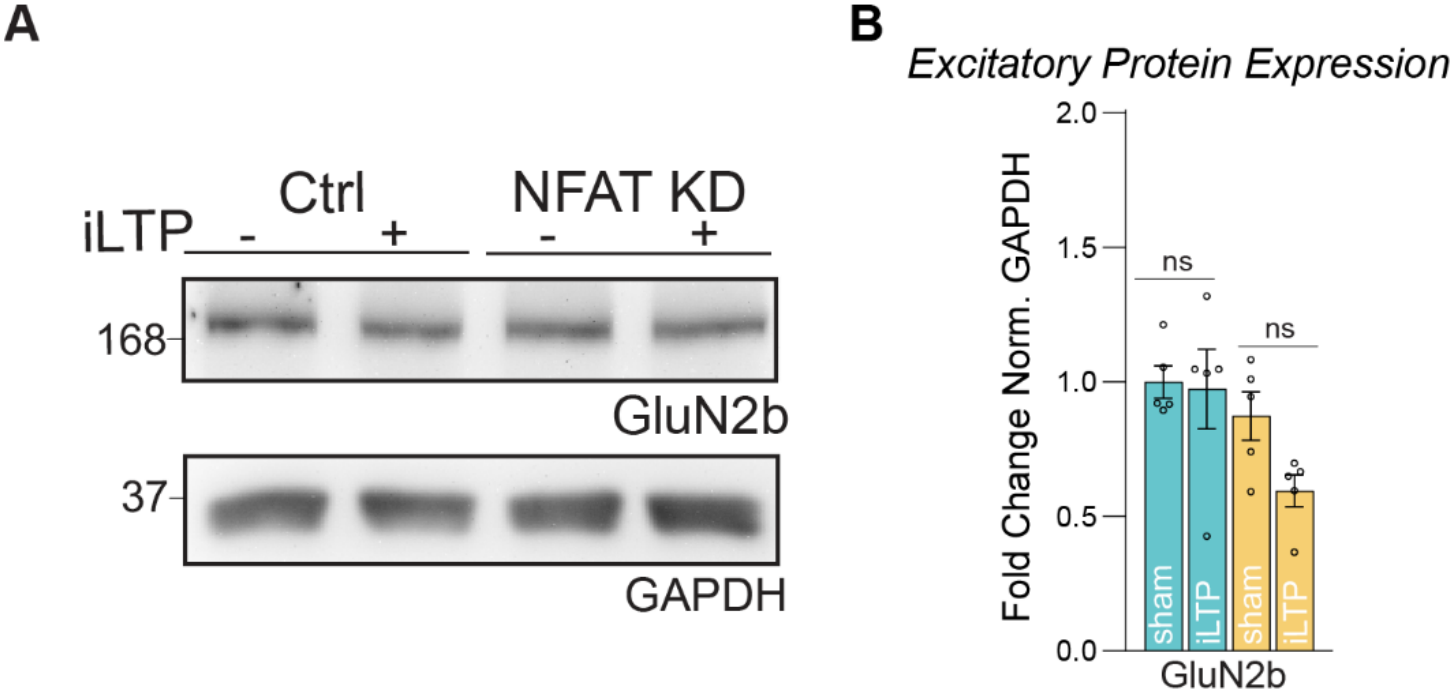
Supplement to Figure 3. NFATc3 KD did not impair excitatory synaptic protein expression. **(A)** Western blots of GluN2b and GAPDH levels in neurons expressing Ctrl (left) or NFATc3 KD (right) either following sham treatment or at 90 minutes post-iLTP. **(B)** Quantification of gephyrin, GluN2b levels in Ctrl or NFATc3 KD neurons. Protein levels were normalized to GAPDH, and data was quantified as relative change in normalized protein expression. n = 5 cultures. (main effect of treatment, F(1, 16) = 2.510, p = 0.1327; main effect of condition, F(1, 16) = 6.885, p = 0.0184; interaction, F(1, 16) = 1.724, p = 0.2077; 2-way ANOVA with Tukey’s multiple comparisons post-hoc test). N = # independent experiments/cultures; n = # neurons. All values represent mean ± SEM. * p < 0.05; 2-way ANOVA with Tukey’s multiple comparisons post-hoc test **(B)**

**Figure S4.**
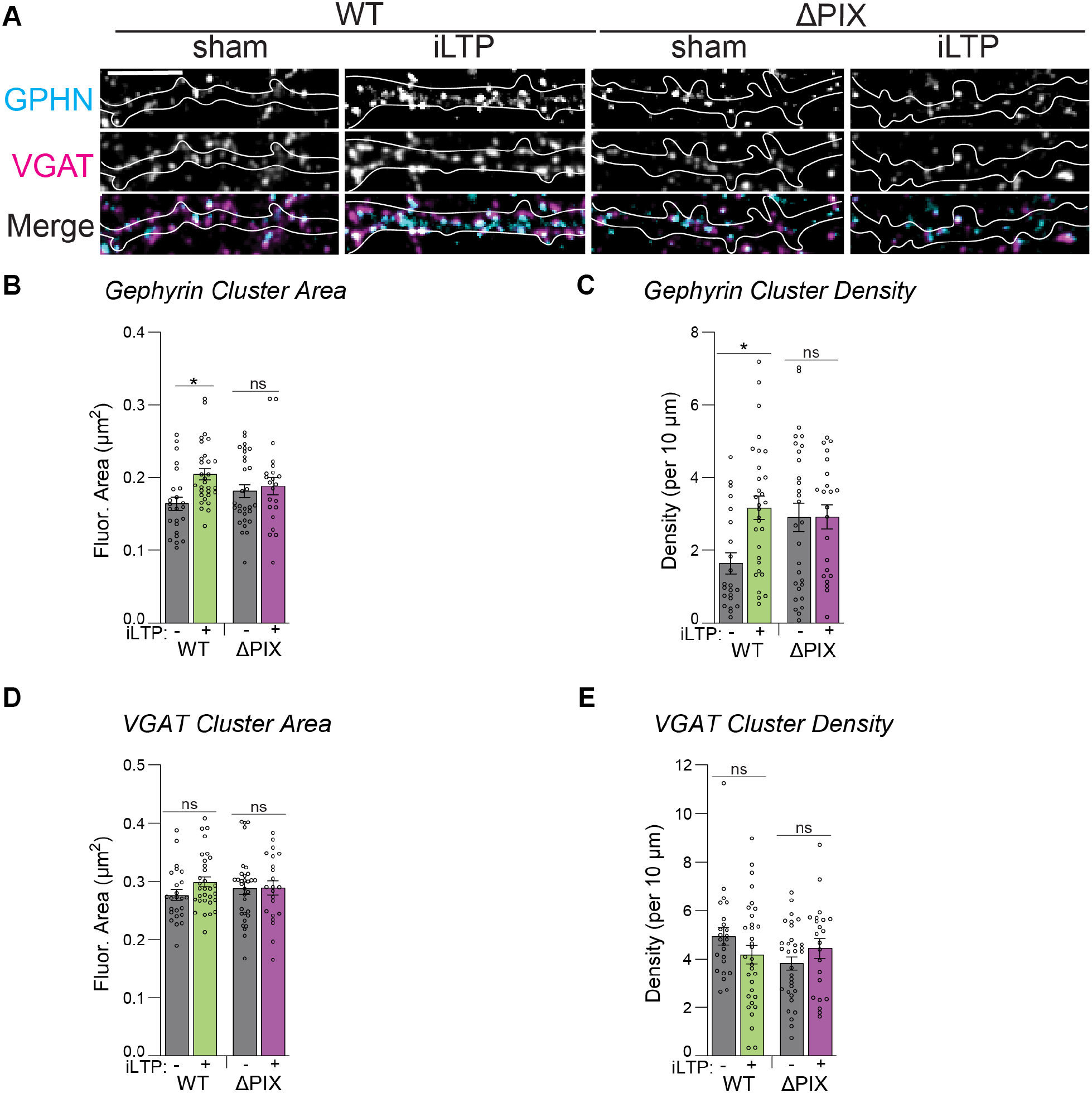
Supplement to Figure 6. AKAP150 anchoring of CaN is required for structural plasticity following iLTP. **(A)** Representative dendritic segments of transfected rat hippocampal neurons expressing AKAP150shRNA + mCh-AKAP79WT (WT) or mCh-AKAP79ΔPIX (ΔPIX) in sham or 90 min post-iLTP. Neurons are labeled with antibodies to gephyrin and VGAT. Scale bar = 10 µm. **(B)** Quantification of gephyrin cluster area from **(A)**. N = 3/ n = 22-30 neurons per condition. (main effect of treatment, F(1, 101) = 6.387, p = 0.0130; main effect of condition, F(1, 101) = 0.003428, p = 0.9534; interaction, F(1, 101) = 3.338, p = 0.0706; 2-way ANOVA with Tukey’s post-hoc test). **(C)** Quantification of gephyrin cluster density from **(A)**. N = 3/ n = 22-30 neurons per condition. (main effect of treatment, F(1, 101) = 4.897, p = 0.0292; main effect of condition, F(1, 101) = 2.134, p = 0.1472; interaction, F(1, 101) = 4.783, p = 0.0311; 2-way ANOVA with Tukey’s post-hoc test). **(D)** Quantification of VGAT cluster area from **(A)**. N = 3/ n = 22-30 neurons per condition. (main effect of treatment, F(1, 101) = 1.347, p = 0.2485; main effect of condition, F(1, 101) = 0.003095, p = 0.9557; interaction, F(1, 101) = 1.127, p = 0.2908; 2-way ANOVA with Tukey’s post-hoc test). **(E)** Quantification of VGAT cluster density from **(A)**. N = 3/ n = 22-30 neurons per condition. (main effect of treatment, F(1, 101) = 0.03641, p = 0.8490; main effect of condition, F(1, 101) = 1.389, p = 0.2412; interaction, F(1, 101) = 3.651, p = 0.0587; 2-way ANOVA with Tukey’s post-hoc test). N = # independent experiments/cultures; n = # neurons. All values represent mean ± SEM. * p < 0.05; 2-way ANOVA with Tukey’s post-hoc test **(B-E)**.

**Figure S5.**
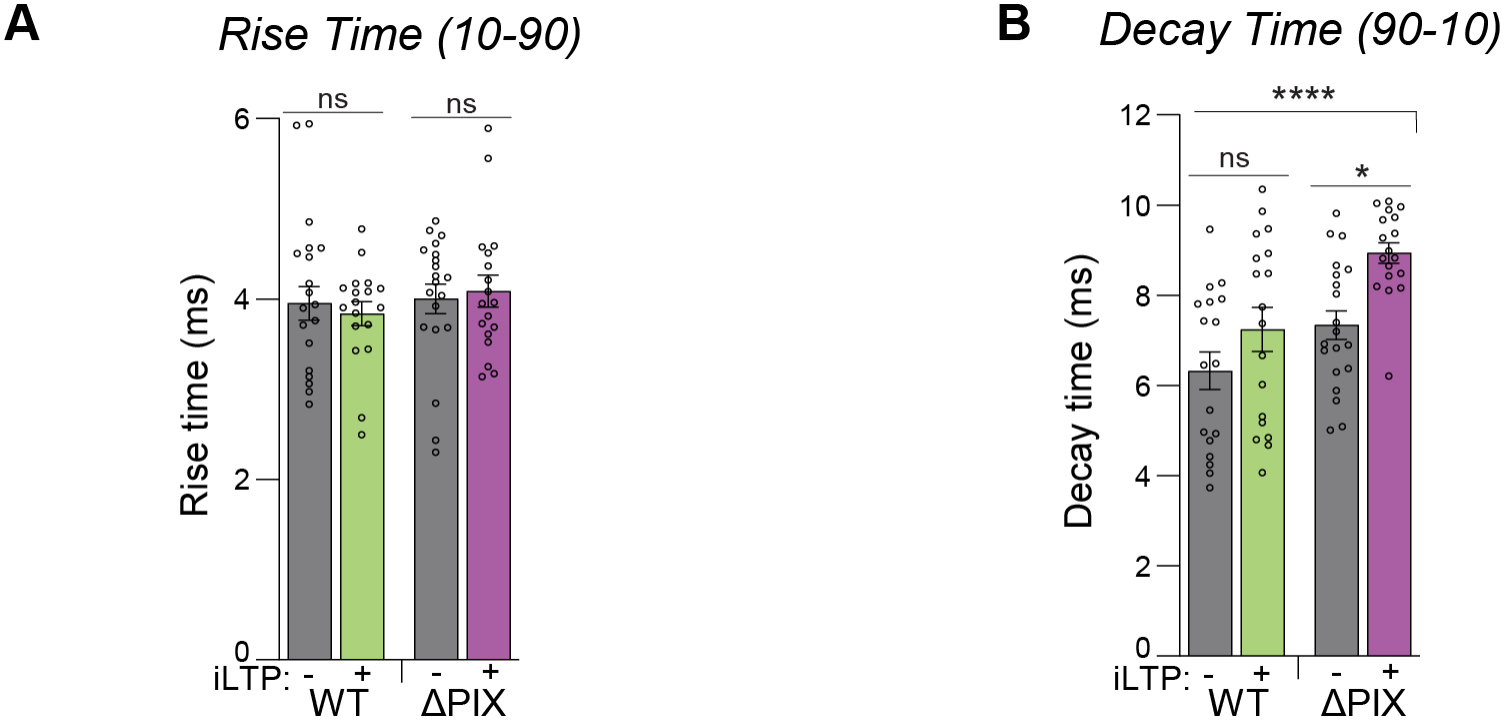
Supplement to Figure 7. Perturbing AKAP-CaN binding increases mIPSC decay time post-iLTP. **(A)** Quantification of mIPSC rise time from WT and ΔPIX neurons in sham or iLTP conditions. N = 5/ n = 18-20 neurons per condition. (main effect of treatment, F(1, 70) = 0.008008, p = 0.9289; main effect of condition, F(1, 70) = 0.8146, p = 0.3699; interaction, F(1, 70) = 0.3561, p = 0.5526; 2-way ANOVA with Tukey’s post-hoc test). **(B)** Quantification of mIPSC decay time from WT and ΔPIX neurons in sham or iLTP conditions. N = 5/ n = 18-20 neurons per condition. (main effect of treatment, F(1, 70) = 11.39, p = 0.0012; main effect of condition, F(1, 70) = 13.13, p = 0.0005; interaction, F(1, 70) = 0.8273, p = 0.3662; 2-way ANOVA with Tukey’s post-hoc test). N = # independent experiments/cultures; n = # neurons. All values represent mean ± SEM. * p < 0.05, **** p < 0.001; 2-way ANOVA with Tukey’s post-hoc test **(A-B)**.

## MATERIALS & METHODS

### Animals

All animal procedures were conducted in accordance with the National Institutes of Health (NIH) United States Public Service guidelines and with the approval of the University of Colorado Denver Institutional Animal Care and Use Committee (Protocol 0426).

### Dissociated hippocampal neurons

Rat hippocampal neurons were dissected from P1-2 mixed sex rat litters and prepared as previously described (24). Hippocampi were dissociated in papain (Worthington), and neurons were plated at a density of 150,000 on 18 mm glass coverslips in 12-well dishes and 1,000,000 in 6-well plates coated with poly-D-lysine (PDL) (Sigma) in Minimum Essential Medium (MEM) and 10% fetal bovine serum (FBS) containing penicillin/streptomycin. The following day, MEM was replaced with Neurobasal media (GIBCO) supplemented with B27 (GIBCO) and Glutamax (GIBCO) and cultured at 37°C, 5% CO_2_, for 14- 18 days. Half of the Neurobasal media and anti-mitotics were changed at DIV 8, followed by a subsequent media change at DIV 12.

Mouse hippocampal neurons were dissected from mixed sex mouse litters from either WT or AKAP150ΔPIX mice prepared as previously described (54). Neurons were plated at a density of 200,000 on 18 mm glass coverslips in 12-well dishes and 1,000,000 in 6-well plates coated with PDL (Sigma). Before experimental use, cells were cultured at 37°C, 5% CO_2_, for 14-18 days in Neurobasal media (GIBCO) supplemented with B27 (GIBCO) and Glutamax (GIBCO). Half of the Neurobasal media and anti-mitotics were changed at DIV 8, followed by a subsequent media change at DIV 12.

### DNA constructs

pmU6-CtrlshRNA-hSyn-mCh (Ctrl), pmU6-NFATc3shRNA-hSyn-mCh (NFATc3 KD), and pmU6-NFATc3shRNA-hSyn-mCh (Resc.) were generated as previously described (23). GFP-NFATc3, GFP-NFATc4, and jRGECO1a were generated as previously described (18). The miRNA upstream regulatory region luciferase constructs were generated as previously described (6, 7). The shRNA KD AKAP150, mCh-AKAP79WT and mCh-AKAP79ΔPIX rescue constructs were generated as previously described (22, 55).

### Lentivirus production

Scrambled (Ctrl) and NFATc3 KD constructs were produced as previously described (23). HEK293T cells (Invitrogen) were cultured at 37°C and 5% CO_2_ in 150 mm dishes. At 60-70% confluence cells were transfected with a Ca^2+^ phosphate-DNA solution containing pCMV-ΔR8.91 packaging plasmid, pCMV-VSV-G envelope plasmid, and pmU6- [shRNA]-hSyn-mCh transfer plasmid (either NFATc3 shRNA or an empty vector control). The following day, HEK cell media was replaced with Neurobasal media (GIBCO) and HEK cells were allowed to produce the virus for 48 hours. Neurobasal culture media was collected by centrifugation of detached HEK cells and supernatant was stored at 4°C for future use.

### Transfections and lentiviral infections

DIV 14-16 mouse and rat hippocampal neurons were transfected with plasmid DNA using Lipofectamine 2000 (Invitrogen) and used for experiments at DIV 16-18. DIV 14-16 rat hippocampal neurons were infected with lentiviral scrambled or NFATc3 KD constructs by applying 250 µL lentiviral supernatants to 6-well dishes and used for experiments at DIV 16-18.

### Drug treatments

iLTP in rat hippocampal neurons was induced by bath application of 20 µM NMDA (Tocris) and 10 µM CNQX (Tocris) in HBS containing (in mM): 145 NaCl, 2 KCl, 10 HEPES, 2 CaCl_2_, 2 MgCl_2_, 10 glucose (pH adjusted to 7.4) in DIV 16-18 rat hippocampal neuronal cultures. iLTP solution was applied at 37°C for 2 minutes as previously described (14). In mouse hippocampal neuronal cultures, iLTP was induced by bath application of 20 µM NMDA in HBS (12, 13) (see above) in DIV 16-18 AKAP150WT or AKAP150ΔPIX cultures. In sham conditions, neurons were treated with a bath application of HBS only for 2 minutes at 37°C.

Cultured neurons were imaged, recorded, fixed or harvested across multiple timepoints following treatment. 5 µM cyclosporin A (CsA) (LC Laboratories) and 5 µM nimodipine (Nim) (Tocris) were added to neuronal cultures 20 minutes prior to iLTP induction and live imaging.

### Whole cell electrophysiology

DIV 16-18 mouse and rat hippocampal neurons expressing mCherry (mCh) were patched in an extracellular artificial cerebrospinal fluid (ACSF) solution containing (in mM): 10 HEPES, 130 NaCl, 5 KCl, 30 D-glucose, 2 CaCl_2_, and 1 MgCl_2_.

Pyramidal neurons were held at -70 mV for 2 minutes prior to recording 3 minutes of continuous neuronal activity with an internal solution containing (in mM): 7.5 CsCl, 67.5 CsMeSO_4_, 0.1 CaCl_2_, 2 MgCl_2_, 10 HEPES, 0.1 EGTA, 0.5 Na_3_GTP, 3 Na_2_ATP, 10 phosphocreatine (290-300 mOsm; pH adjusted to 7.25 with CsOH). mIPSCs were isolated using 1 µM TTX (Tocris), 100 µM APV (Tocris), and 10 µM CNQX (Tocris). mEPSCs were isolated using 1 µM TTX (Tocris) and 30 µM bicuculline (Tocris) and an internal solution containing (in mM): 130 CsMeSO_4_, 3 Na_2_ATP, 0.5 Na_3_GTP, 0.5 EGTA, 10 phosphocreatine, 5 MgCl_2_, 2.5 NaCl, 10 HEPES (290-300 mOsm; pH adjusted to 7.25 with CsOH). Under sham conditions, mIPSCs and mEPSCs were recorded from multiple mCh-expressing pyramidal neurons and the average frequency and amplitude were calculated across recording for sham values. Under iLTP conditions, mIPSCs were recorded 45-90 minutes following treatment and recorded from multiple mCh- expressing pyramidal neurons and the average frequency and amplitude were calculated across the recording for iLTP values.

Data were collected with a Multiclamp 700b amplifier (Molecular Devices) and digitized with a Digidata 1440a (Molecular Devices) at 10 kHz, filtered at 2 kHz, single pole Bessel filter. mIPSCs and mEPSCs were quantified using ClampFit software (Molecular Devices).

### Live imaging

DIV 14-16 rat hippocampal neurons were co-transfected with mCh-shRNA constructs (Ctrl, NFATc3 KD, Resc.) and Gephyrin-FingR-GFP 48-72 hours prior to live imaging. Prior to imaging, coverslips were incubated with VGAT Oyster^650^ (1:200, Synaptic Systems 131 103C5) for 20 minutes at 37°C. Coverslips were mounted in a Ludin chamber (Life Imaging Services) in HBS solution, transfected neurons were imaged pre-stimulation (t = 0 min) and then at various timepoints out to 90 minutes following iLTP stimulation. Confocal images were acquired using an Axio Observer microscope (Zeiss) with a Tokogawa CSU-X1 spinning disk unit, an EC Plan-Neofluar 63x Plan-Apo (1.4 NA) oil immersion objective lens, a Photometrics Evolve 512 EMCCD camera with 16-bit dynamic range, and SlideBook 6.0 software (3i). Images were acquired using 488, 561, and 640 nm laser excitation to capture 3.95 µm z-stacks (13 xy planes, 0.33 µm intervals) in each channel. The maximum intensity of each channel onto 2D images. Gephyrin puncta that were colocalized with VGAT over the 90-minute time course were randomly selected and mean fluorescent intensity was measured following background subtraction at each timepoint in ImageJ. The cluster area and density of gephyrin and VGAT puncta in dendritic segments were analyzed in ImageJ.

### Immunofluorescence microscopy

DIV 16-18 rat hippocampal neurons and WT or AKAP150ΔPIX mouse hippocampal neurons were fixed in 4% paraformaldehyde supplemented with 4% sucrose. Coverslips were incubated in block solution containing: 3% bovine serum albumin (BSA), and 2% normal goat serum (NGS) in PBS for 30 minutes at room temperature. Coverslips were surface stained for GABA_A_R-γ2 (1:500 in block solution, Synaptic Systems 224 004) for 1 hour at room temperature. Following surface staining, coverslips were permeabilized with 0.5% NP-40 for 2 minutes and blocked for 30 minutes prior to total staining of gephyrin (1:750 in block solution, Synaptic Systems 147 111) and VGAT (1:1000 in block solution, Synaptic Systems 131 003) for 1 hour at room temperature. Coverslips were washed 3 x 7 minutes with PBS then incubated with secondary antibodies (1:1000 in block solution, ThermoFisher Alexa-Fluor 488, 568, 647) for 1 hour at room temperature. Coverslips were washed 3 x 12 minutes with PBS then mounted on microscope slides with ProLong gold mounting media (ThermoFisher) and imaged with a spinning disk confocal microscope described in “live imaging” section above. 488, 561, and 640 nm lasers were used to acquire images (3.96 µm stacks) in each channel (13 xy planes, 0.33 µm intervals). The maximum intensity of each channel was projected in a 2D plane and the cluster area and density of each channel were analyzed from dendritic segments using ImageJ.

### Western blotting

DIV 16-18 rat hippocampal neurons and WT or AKAP150ΔPIX mouse hippocampal neurons were harvested in RIPA buffer containing (in mM): 50 Tris (pH 7.5), 1 EDTA, 2 EGTA, 150 NaCl, 1% NP40, 0.5% Na-DOC, 0.1% SDS, 50 NaF, 1 Na_3_VO_4,_ 1:100 dilution from 10x protease inhibitor (Roche). Lysate was harvested in 2x sample buffer containing (in mM): 4% SDS, 20% glycerol, 120 mM Tris at pH 6.8, 0.02% bromophenol blue, 5% 2-mercaptoethanol. Protein samples were boiled for 5 minutes at 95°C and resolved on an SDS-PAGE gel (Bio-Rad). A wet transfer apparatus (Bio-Rad) was used to transfer proteins onto a PDVF membrane (Millipore). Membranes were blocked in 5% milk (Walmart) in TBS-T. Membranes were incubated overnight at 4°C with a primary antibody: GluN2b (1:1000, NeuroMab 75-101), gephyrin (1:10000, Synaptic Systems 147 111), GABA_A_R-α1 (1:500, NeuroMab 75-136), GABA_A_R-β3 (1:5000, NeuroMab 75-149), GABA_A_R-γ2 (1:100, NeuroMab

75-442), and GAPDH (1:10000, GeneTex 627408). Membranes were washed in TBS-T and incubated in HRP-conjugated secondary (1:10,000; Millipore) for 1 hour at room temperature. ECL western blotting substrates (Millipore) were used to visualize protein bands on an Azure 600 imager (Azure Biosystems). Images were analyzed using ImageJ and the protein band of interest was normalized to GAPDH on the same gel.

### RNA Isolation and RT-qPCR

RNA was isolated from rat and WT or AKAP150ΔPIX mouse hippocampal neurons using TRIZOL Reagent (Ambion). RNA extraction was performed by applying chloroform to the samples, mixing vigorously and incubating at room temperature for 3 minutes. Samples were centrifuged at 12,000 g x 15 minutes at 4°C and the aqueous phase was isolated. RNA was precipitated by mixing with ½ volume isopropyl alcohol and incubated for 10 minutes at room temperature. Samples were centrifuged at 12,000 g x 10 minutes at 4°C. Supernatant was removed and the RNA pellet was washed with 75% ethanol and centrifuged at 7,500 g x 5 minutes at 4°C. RNA pellet was air dried for 10 minutes and resuspended in RNase-free water.

mRNAs were reverse transcribed into cDNA using a QuantiTect Reverse Transcription Kit (Qiagen) according to manufacturer’s instructions. cDNA was diluted 1/10 in RNase free water and 1 µL was used per reaction to measure mRNA levels using a QuantiTect SYBR Green PCR Kit (Qiagen). qPCR was performed using gene-specific primers (6, 7) in a CFX384 real time qPCR system (Biorad). All qPCR readings were normalized to GAPDH levels. qPCR cycling parameters were: 95°C 15 min, 94°C 15 sec, 55°C 30 sec, 72°C 30 sec for 40 cycles.

### NFAT translocation imaging

DIV 14-16 rat hippocampal neurons were transfected with GFP-NFATc3 or GFP-NFATc4 constructs 48-72 hours prior to live imaging experiments. To prevent spontaneous network activity, neurons were incubated in tetrodotoxin (TTX; 1 µM) 30 minutes prior to imaging. For drug treatments conditions, coverslips were also treated with either 5 µM cyclosporin A (CsA) (LC Laboratories) or 5 µM nimodipine (Nim) (Tocris) 20 minutes prior to live imaging experiments. Coverslips were mounted in a Ludin chamber (Life Imaging Services) in HBS solution containing TTX, transfected neurons were imaged pre-stimulation (t = 0 min). The iLTP solution in HBS (no TTX) was applied for 2 minutes and HBS with TTX was washed back on following iLTP stimulation. Transfected neurons were imaged at various timepoints out to 90 minutes following iLTP stimulation. Confocal images were acquired with a spinning disk confocal microscope described in “live imaging” section above. A 488 nm laser was used to acquire images (3.96 µm stacks; 13 xy planes, 0.33 µm intervals). The maximum intensity was projected in a 2D plane and the difference between GFP fluorescent intensity was calculated between the nucleus and cytosol using ImageJ then normalized to baseline images. jRGECO1a was imaged using 561 excitation, imaging the baseline, during and immediately following iLTP and ΔF/F_0_ was calculated using soma ROIs normalized to a baseline background ROI.

DIV 14-16 WT or AKAP150ΔPIX mouse hippocampal cultures were transfected with GFP-NFATc3 48-72 hours prior to fixation. Coverslips were incubated in 1 µM TTX for 30 minutes prior to iLTP treatment. The iLTP solution was applied for 2 minutes and conditioned media with TTX was washed back on following iLTP stimulation and neurons were incubated at 37°C and fixed at various timepoints out to 90 minutes following iLTP stimulation. Coverslips were fixed with 4% paraformaldehyde supplemented with 4% sucrose; then mounted on microscope slides ProLong gold mounting media (ThermoFisher) and imaged with a spinning disk confocal microscope described in “live imaging” section above. A 488 laser was used to acquire images (3.96 µm stacks; 13 xy planes, 0.33 µm intervals). The maximum intensity of each was projected in a 2D plane and the difference between GFP fluorescent intensity was calculated between the nucleus and cytosol using ImageJ.

### Luciferase assays

DIV 14-16 WT or AKAP150ΔPIX mouse hippocampal cultures were transfected with 250 ng each of a Firefly luciferase reporter construct and a Renilla luciferase construct. Dual luciferase reporter assays (Promega) were performed according to manufacturer’s instructions. Cultured cells were lysed in a 1x passive lysis buffer for 15 minutes at room temperature on a shaker. Samples were harvested and treated with luciferase assay reagents: Luciferase assay reagent II (LARII) and Stop & Glo reagent (Promega) to measure Firefly and Renilla luciferase activity with a SpectraMax iD3 (Molecular Devices) plate reader luminescence function. Firefly activity was normalized to Renilla activity in each sample.

### Experimental Design and Statistical Analysis

All statistical tests were performed in Prism11 (GraphPad) using t-test, 1-way ANOVA, and 2-way ANOVA with relevant post hoc tests andall data is presented as mean ± SEM. Actual p values are reported in each section of the figure legends. All experiments were performed with 3 separate biological replicates to ensure rigor and reproducibility. Sample size was based on prior studies.

## ACKNOWLEDGEMENTS

This work is supported by NIMH grant R01MH128199 to KRS, NINDS Ruth L. Kirschstein individual predoctoral National Research Service Award (NRSA) F31NS137619 to SEW, NINDS Blueprint Diversity Specialized Predoctoral to Postdoctoral Advancement in Neuroscience (D-SPAN) Award FNS1200640A to JDG, NIMH grant R01MH123700 to MLD.

